# Mutant phenotypes and comprehensive expression analyses reveal roles for CLAVATA in moss vegetative and reproductive development and fertility

**DOI:** 10.1101/2024.04.05.585946

**Authors:** Zoe Nemec Venza, George R. L. Greiff, C. Jill Harrison

## Abstract

The CLAVATA pathway regulates meristem size in angiosperms, but bryophytes have distinct meristematic activities to vascular plants, and gametophytic CLAVATA functions are divergent between species. Here we analysed the promoter activities of all CLAVATA peptide and receptor-encoding genes in the moss *Physcomitrium patens*, and using mutants, identified requirements for PpCLV1 and PpRPK2 receptors in male and female reproductive development and fertility. In gametophytes, all 12 *CLAVATA* genes were expressed in foraging filaments (caulonemata) and leaves (phyllids), but most tissues showed highly specific patterns of promoter activity. *PpCLE3* expression specifically marked gametophyte shoot (gametophore) apical cells and *PpCLV1b* and *PpRPK2* expression marked overlapping apical domains. Expression in male (antheridia) and female (archegonia and eggs) reproductive tissues led us to use mutants to identify roles for *PpCLV1a, PpCLV1b* and *PpRPK2* in fertility and reproductive development. In sporophytes, the foot was a common site of *PpCLE* expression, and all genes were expressed in stomata. No *PpCLE* activity specifically marked the embryonic apical cells, and embryonic *PpCLV1b* and *PpRPK2* expression marked distinct apical and basal domains. Thus, *P. patens* stem cell activity is likely regulated by different genes in gametophytes and sporophytes, and promoter evolution was a likely driver of diversification of CLAVATA function.

**Plain language summary:** Whilst gene gain and duplication contributed to the origin of land plants and diversification of seed plants, significant gene loss is associated with morphological adaptation in bryophytes. In the moss, *Physcomitrium patens*, *CLAVATA* genes expanded from low ancestral numbers, showing exquisite cell type specificity in expression. Our results suggest co-option of CLAVATA into many different developmental processes during moss evolution.

## Introduction

Originally identified with roles in the Arabidopsis fruit and shoot apical meristem (Leyser & Furner, 1992; Clark *et al*., 1993), the CLAVATA pathway is a key short-range signalling mechanism in plants (Fletcher, 2018). CLAVATA pathway genes encode CLE peptides that can diffuse between cells (Fletcher *et al*., 1999; Cock & McCormick, 2001; Rojo *et al*., 2002), and receptor-like kinases such as CLAVATA1 and RECEPTOR-LIKE PROTEIN KINASE2 (RPK2) which perceive the peptides and transduce a signal to downstream components (Clark *et al*., 1997; Trotochaud *et al*., 2000; Kinoshita *et al*., 2010). In the multicellular shoot apex of Arabidopsis, a CLAVATA signalling module comprising the CLAVATA3 (CLV3) peptide, the CLAVATA1 (CLV1) receptor like kinase and the downstream transcription factor, WUSCHEL, operate in a feedback loop to maintain the size and integrity of the meristem as the shoot grows (Mayer *et al*., 1998; Fletcher *et al*., 1999; Brand *et al*., 2000; Schoof *et al*., 2000). Modules with a similar logic but involving different CLEs, receptor-like kinases and *WUSCHEL*-like WOX genes are involved in cell type differentiation and proliferation in roots, the vascular cambium and the expanding leaf margin of Arabidopsis (reviewed in Willoughby & Nimchuk, 2021 and Narasimhan & Simon, 2022). Each gene class comprises many members (Goad *et al*., 2017; Whitewoods *et al*., 2018; Wu *et al*., 2019) which are expressed in diverse cell types and tissues (Jun *et al*., 2010; Fletcher, 2020; Tvorogova *et al*., 2021).

*CLAVATA* genes are land plant specific, and previous studies comparing CLAVATA function between Arabidopsis and moss identified ancestral roles for CLAVATA in stem cell regulation (Whitewoods *et al*., 2018; Cammarata *et al*., 2022). However, mosses and flowering plants have undergone over 470 million years of independent evolution (Morris *et al*., 2018), and divergences in bryophyte and flowering plant CLAVATA function (Sakakibara *et al*., 2014; Hirakawa *et al*., 2019; Hirakawa *et al*., 2020; Cammarata *et al*., 2022) raise questions about the evolution of this short-range signalling mechanism. The moss CLAVATA pathway comprises more components than liverwort or hornwort pathways, suggesting lineage-specific functional diversification (Whitewoods *et al*., 2018; Whitewoods *et al*., 2020). Moss tissues typically have fewer cell layers than flowering plant tissues (Parihar, 1959; Kofuji & Hasebe, 2014). Single celled haploid spores germinate to establish apical cells that cleave to develop filamentous protonemal tissues comprising photosynthetic chloronemata and foraging caulonemata (Parihar, 1959). Both filament types initiate side branches, and in caulonemata, around 5% of these attain bud identity, initiating shoot-like gametophores with leaf-like phyllids (Aoyama *et al*., 2012). Male and female sex organs (antheridia and archegonia) differentiate at the tip of gametophores, respectively producing sperm and eggs (Parihar, 1959). Fertilisation results in diploid zygote formation, embryogenesis and sporophyte development, and sporophyte development terminates in sporangium formation and spore production, completing the life cycle (Parihar, 1959).

The *Physcomitrium patens* (formerly *Physcomitrella patens*) CLAVATA pathway comprises 9 *CLE*s (*PpCLE1-9*) which jointly encode 4 CLV3-like peptides, and 3 receptor like kinases, *PpCLV1a*, *PpCLV1b* (jointly referred to as *PpCLV1*) and *PpRPK2* (Whitewoods *et al*., 2018; Whitewoods *et al*., 2020; Cammarata *et al*., 2022). Previous work identified roles for CLAVATA in protonemal development and gametophore development (Whitewoods *et al*., 2018; Whitewoods *et al*., 2020; Cammarata *et al*., 2022; Nemec-Venza *et al*., 2022). During protonemal development, *PpCLE3*, *PpCLE9* and *PpCLV1a* promoters are active in primary chloronemata, and except *PpCLE1*, *PpCLE8* and *PpCLV1a*, most promoters are active in caulonemal tip cells (Nemec-Venza *et al*., 2022). *PpcleAmiR4-7* and *Pprpk2* knockout mutant protonemata spread further than wild-type plants, and pharmacological experiments and expression analyses in *Ppclv1* and *Pprpk2* mutant backgrounds suggest that *PpCLV1* and *PpRPK2* suppress caulonemal identity by promoting PIN-mediated auxin transport out of filament tip cells (Nemec-Venza *et al*., 2022). Later in the life cycle, *PpcleAmiR4-7*, *Ppclv1a1b* and *Pprpk2* mutants show defects in gametophore initiation, having supernumerary but frequently abortive gametophores (Whitewoods *et al*., 2018; Whitewoods *et al*., 2020). In *Ppclv1b, Ppclv1a1b* and *Pprpk2* mutants, differentiation at the gametophore base is perturbed, and stem cells overproduced (Whitewoods *et al*., 2018; Whitewoods *et al*., 2020; Cammarata *et al*., 2022). Modelling and combinatorial mutant phenotype analysis showed that *PpCLV1* and *PpRPK2* genes act independently via unknown downstream components to suppress gametophore stem cell identity, and that *PpCLV1* or *PpRPK2* represses cytokinin-mediated stem cell initiation (Cammarata *et al*., 2022).

Here we aimed to broaden understanding of CLAVATA function by identifying likely sites of *PpCLE* and receptor-like kinase production throughout the *P. patens* life cycle and examining receptor mutant phenotypes. We found that all tissue types exhibited promoter activity of at least one *PpCLE* and receptor-like kinase gene, with receptor encoding genes typically having broader domains of promoter activity than *PpCLE*s. Whilst all *PpCLE* promoters were active during caulonema and phyllid development, promoter activity was spatiotemporally restricted at other stages, frequently marking a single cell type. As several CLAVATA promoter activities marked specific reproductive cell types, we examined reproductive development in receptor mutants, identifying roles for CLAVATA in fertility, specifically antheridium, archegonium and egg development. In sporophytes, all CLAVATA pathway promoters were active in stomata. *PpCLV1* and *PpRPK2* expression was in mutually exclusive domains during early embryogenesis but later on overlapped in intercalary meristems. Overall, our data identify diverse and likely specific sites of *P. patens* CLAVATA signalling, and potential conservation of CLAVATA function in sporophyte meristems and stomata.

## Results

### PpCLE3, PpCLV1b and PpRPK2 promoters were active in initiating gametophore apices

To resolve the location of CLAVATA activity and likely peptide/receptor like kinase signalling at different lifecycle stages, we first used *promoter::nuclearGFP-GUS* (*pro::NGG*) fusions to characterize patterns of promoter activity at different stages of gametophore development (Figure 1). We defined morphological markers for each stage of bud development; Stage 0 buds had two or three cells, Stage 1 buds had 4-8 cells and a wedge-shaped apical cell, Stage 2 buds had tetrahedral apical cells, visible phyllid primordia and initiating rhizoids, Stage 3 buds had a pointed shape with two or three phyllid primordia covering the apex and Stage 4 buds had two to three fully expanded leaves (Figure 1). All *promoter::NGG* lines except for *PpCLE1::NGG*, *PpCLE8::NGG* and *PpCLV1a::NGG* lines (Figure S1) accumulated signal in a range of tissues in and/or around initiating buds (Figure 1). At Stage 0, staining was most conspicuous in *PpCLV1b::NGG* and *PpRPK2::NGG* lines, and detected in buds and the filament cells immediately underneath them (Figure 1J’ and O’). *PpCLE9::NGG* activity was also observed at Stage 0 (Figure 1E’). No other lines showed conspicuous staining (Figure 1A, F, K, P, U, Z). At Stage 1, apical staining was observed in *PpCLE3::NGG* (Figure 1G), *PpRPK2::NGG* (Figure 1P’) and some *PpCLE9::NGG* buds (Figure 1F’), while lateral staining was observed in *PpCLV1b::NGG* buds (Figure 1K’). Notably, *PpCLE3::NGG* activity specifically marked the apical cell or the apical cell and close daughters throughout subsequent stages of bud development (Figure 1G-J). Staining was usually absent in *PpCLE2::NGG, PpCLE4::NGG, PpCLE5::NGG* and *PpCLE7::NGG* lines (Figure 1B, L, Q, A’). From Stage 2, *PpCLE5::NGG* actvity was strong and specific to rhizoid precursor cells and rhizoid tips (Figure 1R-T) while *PpCLE2::NGG* (Figure 1C), *PpCLE6::NGG* (Figure 1X-Y), *PpCLE7::NGG* (Figure 1C’), *PpCLE9::NGG* (Figure 1H’) and *PpRPK2::NGG* (Figure 1R’) activity was observed later during rhizoid development. *PpCLE7::NGG* (Figure 1B’-C’) activity was observed in the axillary hairs. From Stage 3, *PpCLE2* promoter activity was undetectable (Figure 1D, E), but *PpCLE4::NGG* (Figure 1N-O), *PpCLE6::NGG* (Figure 1X-Y) and *PpCLE9::NGG* (Figure 1H’-I’) promoter activity was detected in single cells close to the apex, and this pattern was maintained at Stage 4. *PpCLV1b::NGG* (Figure 1L’-N’) and *PpRPK2::NGG* (Figure 1Q’-S’) activity marked phyllid primordia and the whole apical region from Stage 2 to Stage 4. Taken with mutant phenotype analyses (Whitewoods *et al*., 2018; Whitewoods *et al*., 2020; Cammarata *et al*., 2022), these distinct patterns of peptide and receptor-encoding gene promoter activity suggest a potential role for short range signaling in the establishment of the apical basal bud axis, the orientation of division planes in the bud and gametophore patterning. We postulate that *PpCLE6, PpCLE9*, *PpCLV1b* and *PpRPK2* are the most likely candidates to control Stage 0 bud cell division planes, and that *PpCLE3* is likely to act with *PpCLV1b* and *PpRPK2* to regulate apical stem function and/or regulate early phyllid development.

**Figure 1:**
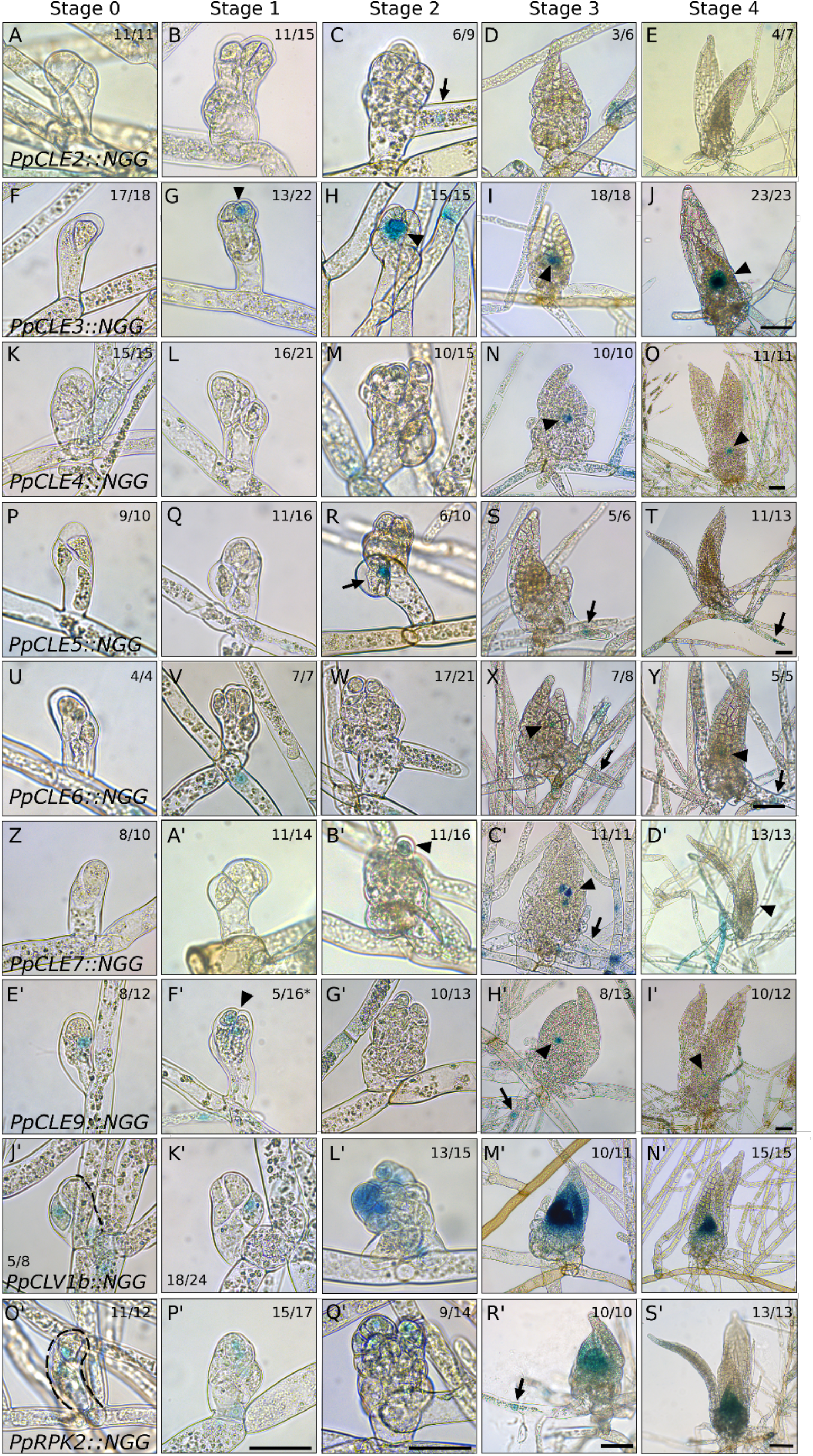
*CLAVATA* promoters were active during gametophore initiation. **(A-E)** *PpCLE2::NGG* activity at (A) Stage 0, (B) Stage 1, (C) Stage 2, (D) Stage 3 and (E) Stage 4 of gametophore development. *PpCLE2::NGG* activity was evident in rhizoids from Stage 2 of development. **(F-J)** *PpCLE3::NGG* activity at (F) Stage 0, (G) Stage 1, (H) Stage 2, (I) Stage 3 and (J) Stage 4 of gametophore development. Activity was undetectable at Stage 0 but marked apical cells from Stage 1 and throughout subsequent development. **(K-O)** *PpCLE4::NGG* activity at (K) Stage 0, (L) Stage 1, (M) Stage 2, (N) Stage 3 and (O) Stage 4 of gametophore development. Activity was detected in apical cells from Stage 3 of development onwards. **(P-T)** *PpCLE5::NGG* activity at (P) Stage 0, (Q) Stage 1, (R) Stage 2, (S) Stage 3 and (T) Stage 4 of gametophore development. Activity marked rhizoid initiation and subsequent development. **(U-Y)** *PpCLE6::NGG* activity at (U) Stage 0, (V) Stage 1, (W) Stage 2, (X) Stage 3 and (Y) Stage 4 of gametophore development. *PpCLE6* promoter activity was detected in cells subtending Stage 0 and Stage 1 buds, and inthe apical region and/or rhizoids from Stage 3 of gametophore development. **(Z-D’)** *PpCLE7::NGG* activity at (Z) Stage 0, (A’) Stage 1, (B’) Stage 2, (C’) Stage 3 and (D’) Stage 4 of gametophore development. *PpCLE7::NGG* expression was detected in apical hairs and rhizoids. **(E’-I’)** *PpCLE9::NGG* activity at (E’) Stage 0, (F’) Stage 1, (G’) Stage 2, (H’) Stage 3 and (I’) Stage 4 of gametophore development. Activity was detected in 2-celled buds, the apical portion of a minority of Stage 1 buds, and rhizoids and phyllid initials in Stage 3 buds. **(J’-N’)** *PpCLV1b::NGG* activity at (J’) Stage 0, (K’) Stage 1, (L’) Stage 2, (M’) Stage 3 and (N’) Stage 4 of gametophore development. Activity was detected at the base of 2- or 4-celled buds but was stronger in phyllid primordia and the apex from Stage 2. **(O’-S’)** *PpRPK2::NGG* activity at (O’) Stage 0, (P’) Stage 1, (Q’) Stage 2, (R’) Stage 3 and (S’) Stage 4 of gametophore development. *PpRPK2* promoter activity was detected throughout buds and was later detected in apices, phyllid primordia and rhizoids. Arrowheads indicate expression in the apical region, arrows indicate rhizoid expression. Numbers indicate the proportion of buds with a similar expression pattern. * indicates a minority of buds were stained. Scale bars Stage 0-2 = 50 µm, Scale bars Stage 3-4 = 100 µm.

### All promoters were active in developing gametophores

To further investigate roles for CLAVATA in gametophore and phyllid development, we stained and imaged gametophores from 4 week-old plants (Figure 2). Whilst *PpCLE3*, *PpCLE4*, *PpCLE6, PpCLE7* and *PpCLE9* promoters were persistently active in gametophore apices (Figure 2A), *PpCLE3::NGG* activity appeared specific to apical cells and branch meristems initiating at the gametophore base whereas *PpCLE9::NGG* activity was broader (Figure 2A). *PpCLV1b::NGG* and *PpRPK2::NGG* also showed activity in the apical region (Figure 2A). *PpCLE6::NGG*, *PpCLV1a::NGG*, *PpCLV1b::NGG* and *PpRPK2::NGG* showed activity in the gametophore axis (Figure 2A), and *PpCLE5::NGG* activity specifically marked rhizoid development as in buds (Figure 2A). To some extent all promoters were active during phyllid development with the stage varying by promoter and most promoters showing activity at the phyllid base throughout development (Figure 2B). In expanding phyllids (P1-P4, to the left of the dashed line), *PpCLE6::NGG* activity was observed at the midrib base, *PpCLE9::NGG* activity marked the midrib and *PpCLV1b::NGG* and *PpRPK2::NGG* activity marked the base in both the midrib and the lamina (Figure 2B). As or after phyllids attained their maximum length (P5 or older, to the right of the dashed line), a wave of promoter activity proceeding from phyllid tips to bases was observed in *PpCLE1::NGG*, *PpCLE2::NGG*, *PpCLE7::NGG* and *PpRPK2::NGG* lines (Figure 2B). Similar waves of promoter activity started from the middle portion of the lamina in *PpCLV1a::NGG* and *PpCLV1b::NGG* lines and were observed in the midrib of *PpCLE1::NGG* and *PpCLV1a::NGG* lines (Figure 2B). *PpCLV1a::NGG*, *PpCLV1b::NGG* and *PpRPK2::NGG* activity persisted at the base of fully developed phyllids (Figure 2B). *PpCLE3::NGG*, *PpCLE4::NGG*, *PpCLE5::NGG* and *PpCLE8::NGG* activity was largely specific to juvenile phyllids (far right of Figure 2B). Thus, CLAVATA promoter activity was observed at diverse developmental stages in a range of gametophore tissues, with varying degrees of cell type specificity. Our results indicate *PpCLE3* promoter activity as an apical cell marker, *PpCLE5* promoter activity as a rhizoid marker and *PpCLE9* promoter activity as a midrib marker.

**Figure 2:**
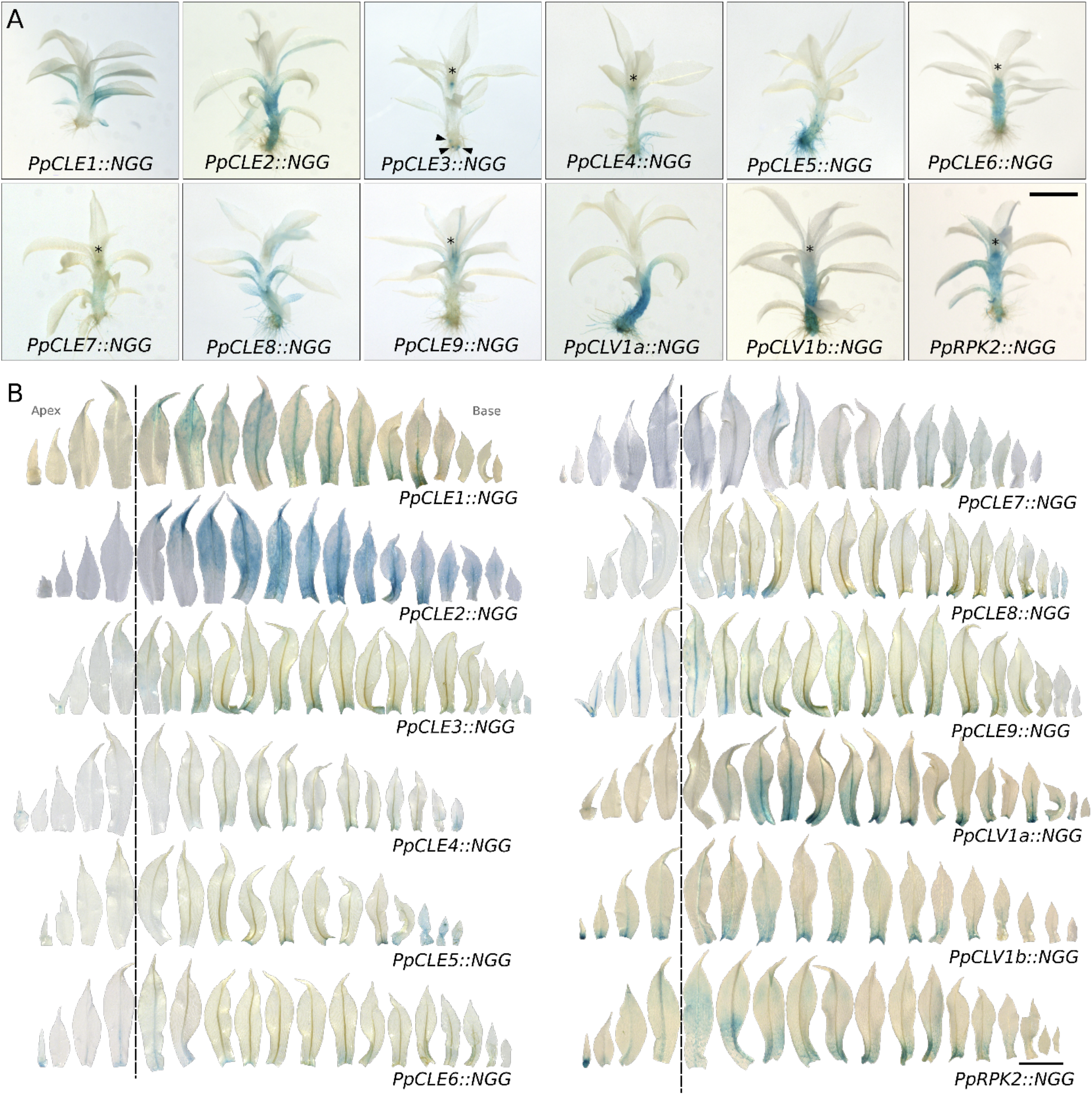
CLAVATA promoters were active during gametophore development. **A)** In gametophores, *PpCLE3::NGG*, *PpCLE4::NGG*, *PpCLE6::NGG*, *PpCLE7::NGG*, *PpCLE9::NGG, PpCLV1b::NGG* and *PpRPK2::NGG* lines showed activity around the apical region (asterisk), and signal in *PpCLE3::NGG* lines was also observed in the apical cells of branches initiating at the gametophore base (arrowheads). *PpCLE2::NGG*, *PpCLE6::NGG*, *PpCLV1a::NGG*, *PpCLV1b::NGG* and *PpRPK2::NGG* promoter activity was detected in the gametophore axis, and *PpCLE5::NGG* was active in rhizoids from early emergence. Phyllid staining was observed in all lines. **B)** At early stages of phyllid development (P1-P4 to the left of the dashed lines) *PpCLE6::NGG* activity was detected in the basal portion of the midrib, and *PpCLE9::NGG* was active in the whole midrib. *PpCLV1b::NGG* and *PpRPK2::NGG* activity was detected in the basal portion of the lamina and midrib. In fully expanded phylllids (P5 or more to the right of the dashed lines), basipetal waves *of PpCLE1::NGG*, *PpCLE2::NGG*, *PpCLE3::NGG*, *PpCLE7::NGG*, *PpCLV1a::NGG*, *PpCLV1b::NGG* and *PpRPK2::NGG* activity were detected. At least three phyllid series were imaged from each line with replicates showing similar results. Scale bars = 1 mm

### PpCLV1a, PpCLV1B and PpRPK2 regulate fertility

Patterns of promoter activity shown in Figures 1 and 2 help to account for defective bud and gametophore development mutant phenotypes previously reported by (Whitewoods *et al*., 2018; Whitewoods *et al*., 2020; Cammarata *et al*., 2022). However, previous work did not investigate roles for the *P. patens* CLAVATA pathway in reproductive development. During the reproductive phase transition, archegonia (egg-producing female gametangia) and antheridia (sperm-producing male gametangia) are produced at the shoot apices. Moisture triggers the rupture of the sac-like antheridium and sperm cell release, and sperm are attracted to mature archegonia. The basal cavity of each archegonium contains an egg cell, and sperm cells swim through the hollow archegonium neck to reach the egg. Fertilisation results in diploid zygote formation, and this is followed by embryo and sporophyte development (Parihar, 1959). Our first investigation into potential roles for CLAVATA in reproductive development comprised a fertility analysis in wild-type versus receptor mutant plants (Figure 3). This showed that *Ppclv1a1b*, *Pprpk2* and *Ppclv1a1brpk2* mutant plants were sterile (Figure 3A). While 43% of wild-type gametophores produced sporophytes, only 16% of *Ppclv1a* and 1.5% of *Ppclv1b* gametophores produced sporophytes (Figure 3B). Thus, *PpCLV1a and PpCLV1b* are jointly required for sporophyte development, with *PpCLV1b* playing a greater role. *PpRPK2* may act independently or together with *PpCLV1* to regulate fertility.

**Figure 3:**
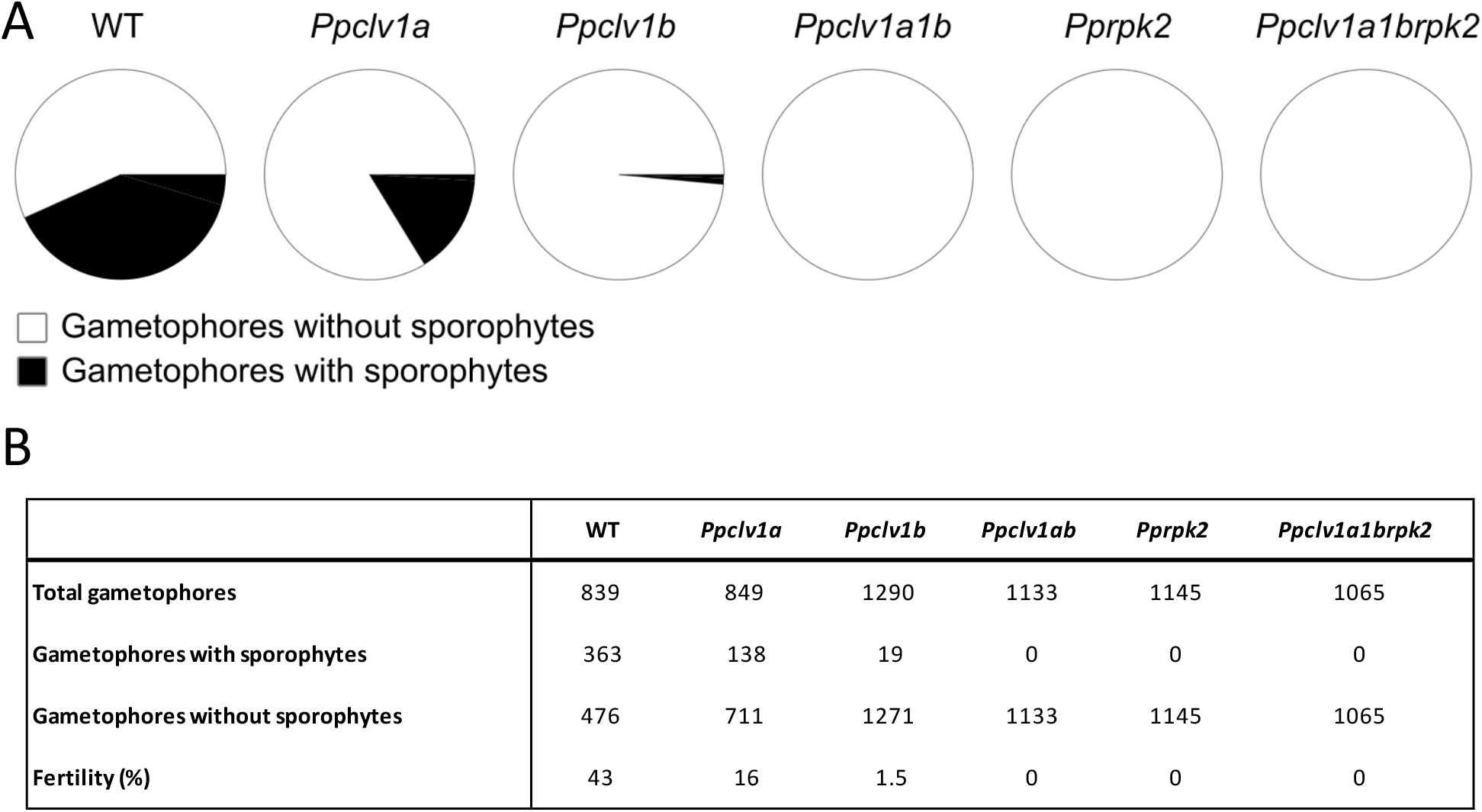
Fertility impairment in CLAVATA mutants. **A)** Pie charts comparing the proportion of gametophores with or without sporophytes in wild-type versus mutant plants after growth in inductive conditions. **B)** Table showing frequency of sporophyte development in wild-type and mutant plants. Pooled data from at least two experimental replicates were used.

### PpCLE6, PpCLE9, PpCLV1a and PpRPK2 promoters were active in antheridia

To identify the developmental basis of the fertility defects above, we next analysed patterns of peptide and receptor gene promoter activity and receptor mutant phenotypes in developing gametangia, starting with antheridia (Figure 4). Antheridial development was previously staged by Landberg *et al*. (2013), who identified formative cell divisions during Stages 1-5 of development and sperm cell maturation from Stage 6 to 9 of development. Sperm cells are initially round (Stages 6 and 7) before elongating to become filiform (Stages 8 and 9), and sperm cells are released as antheridia open at Stage 9; an empty antheridial sac comprises Stage 10 (Landberg *et al*., 2013). While *PpCLE2::NGG*, *PpCLE7::NGG* and *PpCLV1b::NGG* promoter activity was undetectable during antheridium development, *PpCLE1::NGG*, *PpCLE3::NGG, PpCLE4::NGG*, *PpCLE5::NGG* and *PpCLE8::NGG* lines showed weak staining in a minority of antheridia at Stages 6 and 7 (Figure S2). The *PpCLE6* promoter was active at all developmental stages showing strong staining in the antheridial stalks (Figure 4A). Conversely, the *PpCLE9* promoter was active in the apical portion of antheridia, with more intense staining in the apical cell that later ruptures to allow antheridial opening. Basal *PpCLV1a::NGG* activity was detected up to Stage 7 of antheridium development, and activity was later observed throughout a minority of Stage 9 samples (Figure 4A). *PpRPK2::NGG* activity was observed throughout the Stage 1-5 antheridium, with less intensity at Stage 6/7 and then more intensity at Stage 9/10 (Figure 4A). Overall, these staining patterns suggest the presence of apical (*PpCLE9*) and basal (*PpCLE6*) sources of CLE peptides during antheridium development, with likely perception by PpCLV1a and/or PpRPK2 receptors.

**Figure 4:**
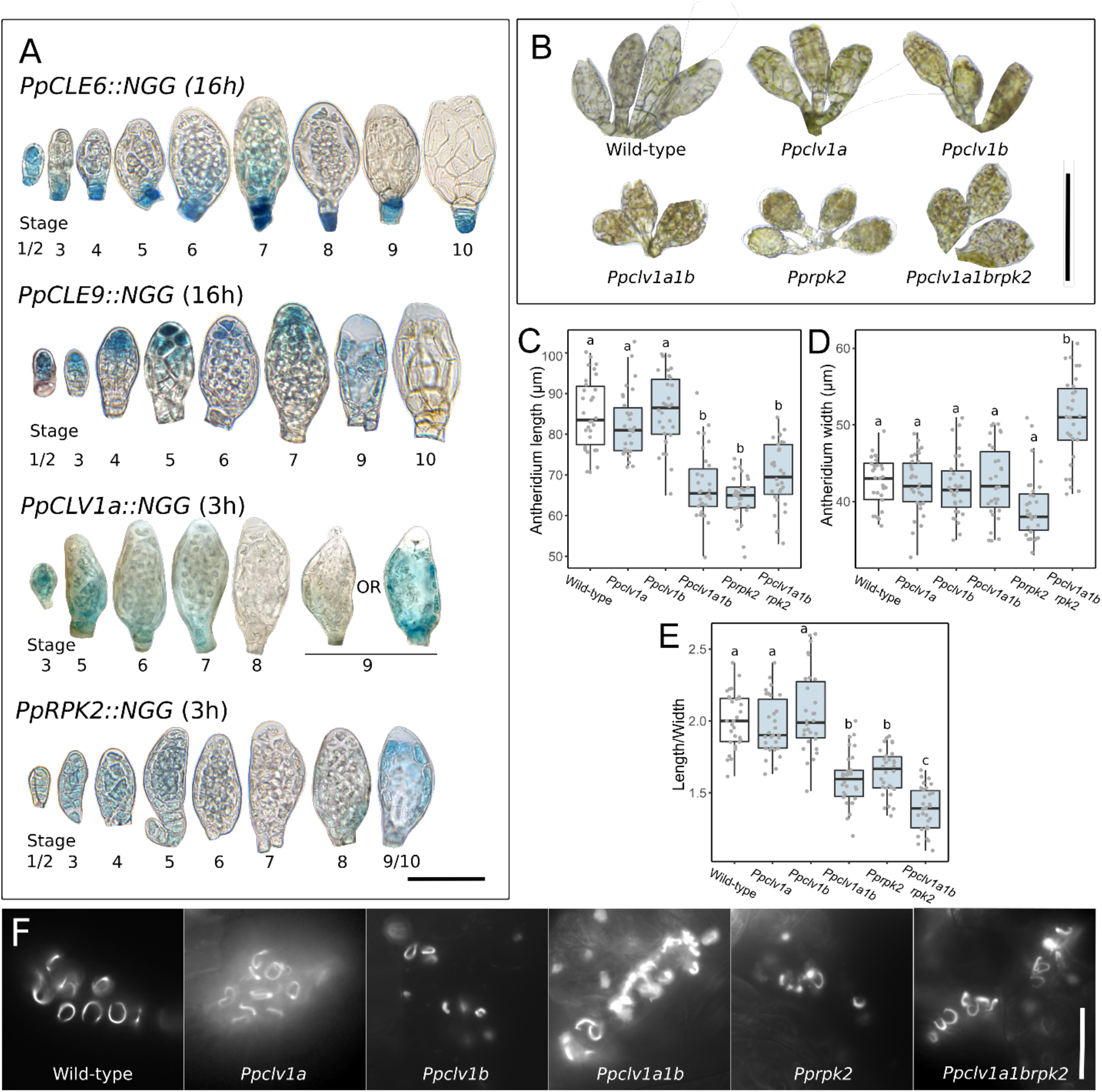
CLAVATA regulates male reproductive development. **A)** Light micrographs of GUS stained and dissected antheridia showing *PpCLE6::NGG*, *PpCLE9::NGG, PpCLV1a::NGG* and *PpRPK2::NGG* activity at different stages of development. Whilst *PpCLE6::NGG* and *PpCLV1a::NGG* activity was manly basal, *PpCLE9::NGG* activity was mainly apical and *PpRPK2::NGG* activity extended throughout developing antheridia. Staining was less intense at Stage 8 and Stages 6 to 8 in *PpCLV1a::NGG* and *PpRPK2::NGG* lines. Only some *PpCLV1a::NGG* antheridia at Stage 9 showed expression. Staining times are displayed in brackets. Scale bar = 50µm. **B**) Light micrographs of antheridia showing developmental defects in *Ppclv1a1b*, *Pprpk2* and *Ppclv1a1brpk2* mutants. Scale bar = 100 µm. **C**) Boxplot showing that *Ppclv1a1b*, *Pprpk2* and *Ppclv1a1brpk2* mutants had shorter antheridia than wild-type and *Ppclv1a* and *Ppclv1b* mutant plants. N >30. **D**) Boxplot showing that *Pprpk2* and *Ppclv1a1brpk2* mutants had respectively narrower or wider antheridia than wild-type plants. N > 30. **E)** Boxplot showing that *Ppclv1a1b, Pprpk2* and *Ppclv1a1brpk2* mutants had more globose antheridia than wild-type plants N > 30. In all graphs, mid lines represent median values, boxes represent the 1st to 3rd quartile range, whiskers represent 95% confidence interval. Different letters indicate significant differences between groups identified by ANOVA and Tukey test. **F**) Fluorescence micrographs of mature DAPI stained sperm from wild-type and receptor mutant plants showing sperm with normal morphology. Scale bar = 20 µm.

### PpCLV1a, PpCLV1b and PpRPK2 regulate male reproductive development

To investigate CLAVATA receptor functions in antheridium development, we induced reproduction in *Ppclv1a*, *Ppclv1b*, *Ppclv1a1b* and *Pprpk2* and *Ppclv1a1brpk2* mutants and compared the morphology of fully developed wild-type and mutant antheridia (Figure 4B). While gross antheridium structure appeared normal in mutants, *Ppclv1a1b, Pprpk2* and *Ppclv1a1brpk2* mutant antheridia were significantly shorter than wild-type antheridia and *Ppclv1a1brpk2* antheridia wider than wild-type antheridia (Figure 4C-D). Mutant antheridia were smaller and more globose than wild-type antheridia (Figure 4E). Although sperm cells with normal morphology were ultimately recovered from all mutant lines, the vast majority of *Pprpk2* antheridia were found empty (Figure 4F). Thus, defects in male reproductive development may contribute to *Pprpk2* mutant fertility defects but are unlikely to underly sterility in *Ppclv1a1b* mutants (Figure 3).

### PpCLE3, PpCLE4, PpCLE9, PpCLV1a, PpCLV1b and PpRPK2 promoters were active in archegonia

To evaluate the potential contribution of CLAVATA to female reproductive development, we next examined the activity of peptide and receptor-encoding gene promoters in archegonia at different stages of development (Figure 5). Archegonia undergo formative cell divisions at Stages 1-5 of development, and cell division and elongation then lengthen the neck at Stages 5–7 (Landberg *et al*., 2013). Cell division ceases at Stage 7, and cells at the apices rupture at Stage 9 to open the hollow neck cavity which turns brown at Stage 10 (Landberg *et al*., 2013). *PpCLE1::NGG*, *PpCLE2::NGG*, *PpCLE5::NGG*, *PpCLE7::NGG* and *PpCLE8::NGG* activity was undetectable or detected in a minority of samples (Figure S2). Whilst *PpCLE6::NGG* activity was weak and diffuse through archegonium development, *PpCLE3::NGG*, *PpCLE4::NGG* and *PpCLE9::NGG* activity was strong and consistent in eggs, canal cells or both, with *PpCLE3::NGG* and *PpCLE9::NGG* activity starting at early developmental stages (Figure 5A). *PpCLE9::NGG* activity was detected in the tip cells of Stage 4 to 7 archegonia, and broad *PpCLE4::NGG* activity in the neck was detected at Stage 7 to 9 of archegonial development (Figure 5A). Broad *PpCLV1a::NGG* activity was detected up to Stage 4 of archegonium development, and later became stronger in the internal egg an canal cell lineage (Figure 5B). *PpCLV1b::NGG* activity was primarily detected in egg cells at Stages 9 and 10 of development (Figure 5B). *PpRPK2::NGG* activity was broad up to Stage 4 of archegonium development, but was stronger at the base from Stages 5-6 and was present in the egg cell at Stages 9 and 10 of development (Figure 5B). Thus, *PpCLE3::NGG*, *PpCLE4::NGG*, *PpCLE6::NGG, PpCLE9::NGG*, *PpCLV1a::NGG*, *PpCLV1b::NGG* and *PpRPK2::NGG* activity indicated potential roles for CLAVATA peptides and receptors in female reproductive development.

**Figure 5:**
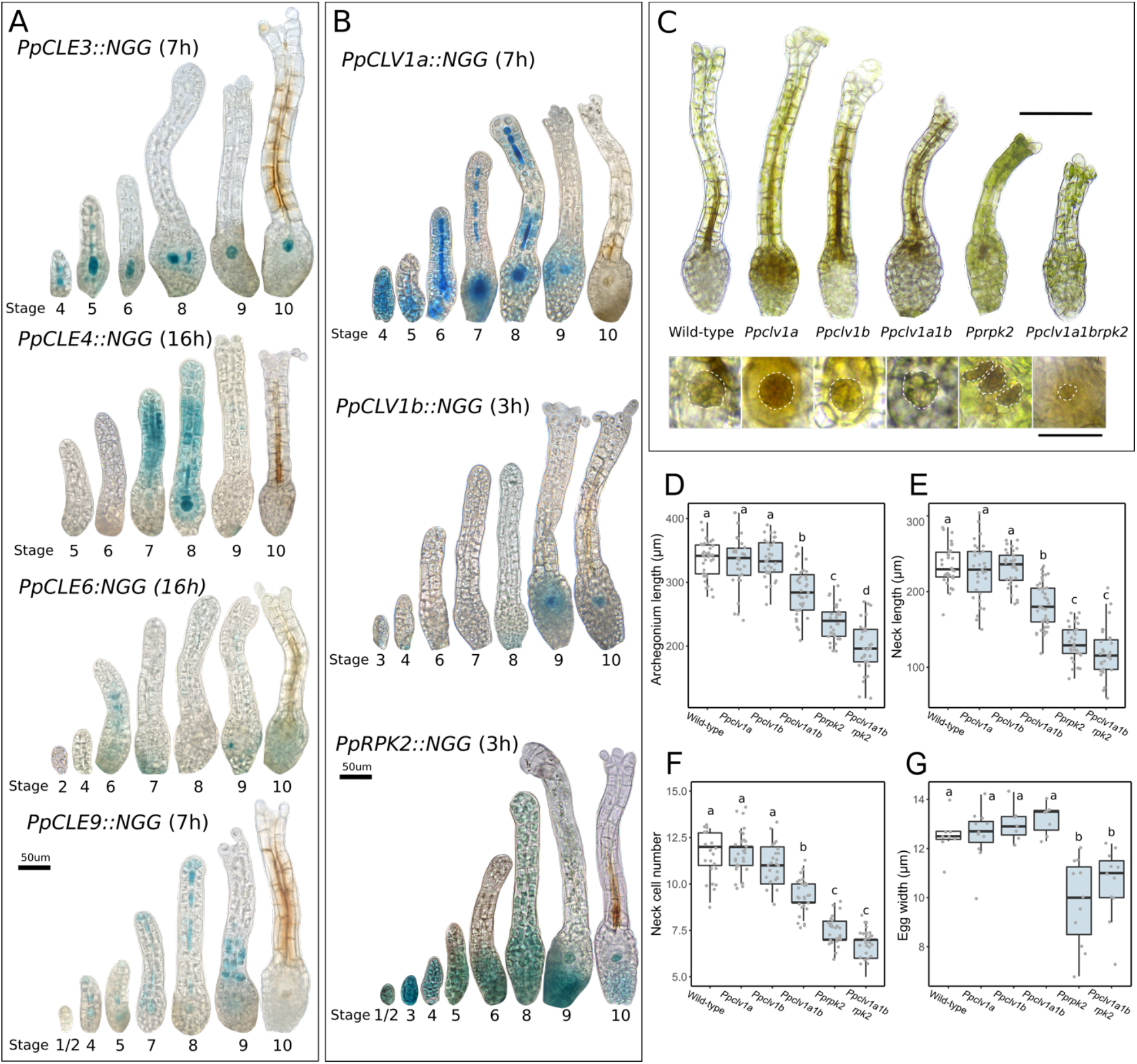
CLAVATA regulates female reproductive development. **A)** Light micrographs of GUS stained archegonia at different stages of development showing peptide-encoding gene promoter activity. *PpCLE3::NGG* activity was detected in inner cells of the neck and the egg cell lineage, *PpCLE4::NGG* activity was detected in the Stage 7 and 8 neck and developing egg and *PpCLE6::NGG* activity was broad and diffuse in venters. *PpCLE9::NGG* activity marked the tip, inner neck cells and egg cell lineage, and after opening staining was present at the base of the neck. **B)** Light micrographs of archegonia at different stages of development showing receptor-encoding gene promoter activity. *PpCLV1a::NGG* activity was detected throughout Stage 4 archegonia, at later stages narrowing down to the inner cells of the neck and developing egg. *PpCLV1b::NGG* activity mainly marked egg cells at Stages 9 and 10 of development. *PpRPK2::NGG* activity was present throughout the initiating archegonium (Stages 1-3), later narrowing to the venter (Stages 5 to 10) and developing eggs. Staining times are displayed in brackets. Scale bar = 50 µm **C)** Light micrographs of fully developed archegonia and eggs (white dashed lines) from wild-type, *Ppclv1a*, *Ppclv1b*, *Ppclv1a1b*, *Pprpk2* and *Pprpk2clv1a1b* mutant plants. Scale bars = 100 μm, 25 µm. **D)** Box plot showing mature archegonium lengths in wild-type and receptor mutant plants; n > 30. **E)** Boxplot showing mature archegonial neck lengths in wild-type and receptor mutant plants; n > 30. **F)** Boxplot showing the number of cells in the archegonial neck in wild-type and receptor mutant plants; n > 22. **G)** Boxplot showing egg width in wild-type and receptor mutant plants; n = 7-11. In all box plots midlines represent the median value, boxes represent the interquartile range, whiskers represent 95% confidence intervals. Different letters indicate significant differences between groups identified by ANOVA and Tukey test.

### PpCLV1a, PpCLV1b and PpRPK2 regulate female reproductive development

To further investigate such potential roles we next analysed receptor mutant phenotypes (Figure 5C-G). While *Ppclv1a* and *Ppclv1b* mutant archegonia resembled those of wild-type plants, and archegonial initiation and overall structure appeared normal in all mutants, archegonial morphology was perturbed in *Ppclv1a1b*, *Pprpk2* and *Ppclv1a1brpk2* mutants. Mutant archegonia had shorter necks with fewer cells than wild-type archegonia (Figure 5D-F), and *Ppclv1a1brpk2* mutants also had reduced overall length (Figure 5D), showing an additive effect of *Ppclv1a1b* and *Pprpk2* mutations. Mutants also displayed defects in egg development (Figure 5C). Whereas wild-type, *Ppclv1a*, *Ppclv1b* and *Ppclv1a1b* mutant eggs were similar sizes and spherical, *Pprpk2* and *Ppclv1a1brpk2* mutant egg development was aberrant, resulting in the formation of several smaller cells with irregular shapes and sizes (Figure 5C, G). Thus, defects in egg development may underly *Pprpk2* mutant sterility, but *Ppclv1a1b* mutant sterility (Figure 3) had no obvious basis in female reproductive development.

### Promoter activities indicate diverse roles for CLE peptides and their receptors in sporophyte development

As *Ppclv1a1b*, *Pprpk2* and *Ppclv1a1brpk2* mutants were sterile, it was not possible to identify roles for CLAVATA in sporophyte development by mutant phenotype analysis. However, we were able to identify likely sites of CLAVATA peptide and receptor production using *promoter::NGG* lines (Figure 6). During sporophyte development, the apical cells first produce lateral merophytes (Stage 1) which then start dividing to make the embryonic axis (Stage 2) (Coudert *et al*., 2019). The apical cell arrests (Stage 3) and an intercalary meristem around the middle of the embryo activates to extend the embryonic axis (Stage 4) (Coudert *et al*., 2019). The apical portion of the sporophyte then swells to form the sporangium (Stage 5) which contains the spore mass and columella and has stomata in a ring around the base (Stage 6) (Coudert *et al*., 2019). The capsule becomes spherical at Stage 7 (Coudert *et al*., 2019). At early developmental Stages (1-3), *PpCLE3::NGG*, *PpCLE6::NGG* and *PpCLE9::NGG* activities were broad, and *PpCLE2::NGG*, *PpCLE3:NGG*, *PpCLE6:NGG*, *PpCLE7::NGG*, *PpCLE9::NGG* lines showed activity in parental tissue below the sporophyte foot (Figure 6B, C, F, G, I). *PpCLE7::NGG* and *PpCLE8::NGG* were active in the hypophysis from Stage 4 (Figure 6 G, H) and all nine *PpCLE::NGG* reporters were active in stomata (Figure 6A-I). *PpCLE1::NGG*, *PpCLE2::NGG*, *PpCLE3::NGG*, *PpCLE6::NGG* and *PpCLE9::NGG* activity was detected in the sporophyte foot (Figure 6A-C, F, I). Some patterns of promoter activity were gene-specific, for instance, *PpCLE4::NGG* activity was detected in sporogenic tissue (Figure 6D), while *PpCLE2::NGG* activity was detected in a few cells at the upper end of the developing spore mass and *PpCLE6::NGG* activity was detected at the lower end (Figure 6B, F). *PpCLE3::NGG*, *PpCLE6::NGG* and *PpCLE9::NGG* activity was detected in groups of cells in the intercalary region, and *PpCLE9::NGG* activity was in apical-basal cell files where water-conducting cells later differentiate (Figure 6C, F, I). Unlike *PpCLE* promoter activity, and their activity in gametophytes, *PpCLV::NGG* and *PpRPK2::NGG* activity overlapped little in sporophytes (Figure 6J-L). In *PpCLV1a::NGG* lines, sporophyte development could only be observed to Stage 3 or 4 due to a background phenotypic defect. However, *PpCLV1a::NGG* activity was detected in sporogenic tissues in a similar pattern to *PpCLE4::NGG* activity (Figure 6J). While *PpCLV1b::NGG* activity was strong in the apical half of the sporophyte (Figure 6K), *PpRPK2::NGG* activity was strong in the basal sporophyte and later formed a band in the stomatal region (Figure 6L). *PpCLV1b::NGG* and *PpRPK2::NGG* activity appeared juxtaposed or overlapping at Stages 3-5 when the intercalary meristem is active. Overall, promoter activities suggest production of multiple CLE peptides in stomata and at the sporophyte foot and, their likely perception by *PpCLV1b* in the apical and *PpRPK2* in the basal sporophyte.

**Figure 6:**
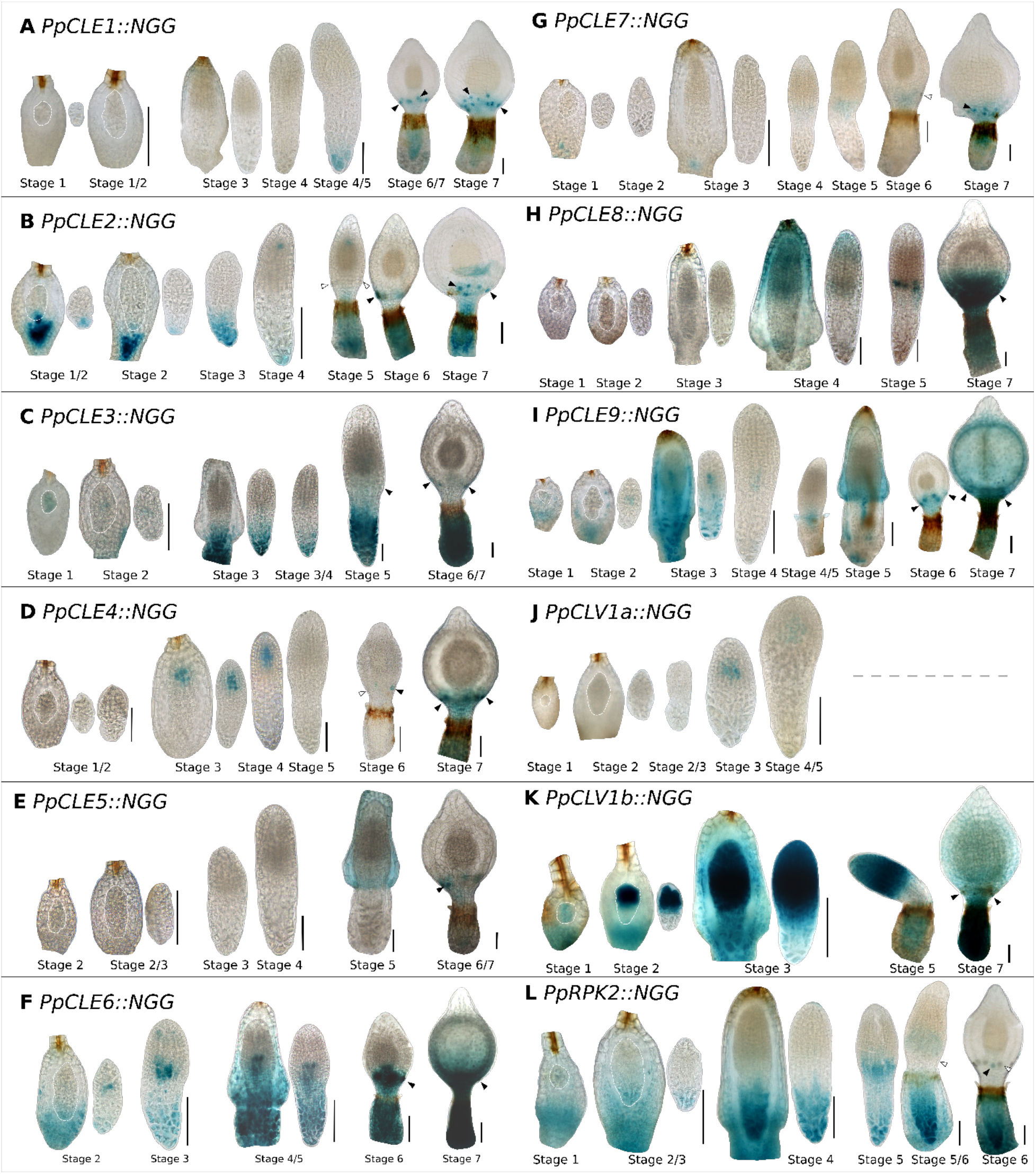
*PpCLE*, *PpCLV1a*, *PpCV1b* and *PpRPK2* promoters were active in diverse sporophytic tissues. **A)** *PpCLE1::NGG* signal was detected at the sporophyte foot and in stomata. **B)** *PpCLE2::NGG* signal was detected in maternal tissue, and in the sporophyte foot, stomata and a few apical sporogenic cells. **C)** *PpCLE3::NGG* signal was detected in maternal tissue, Stage 1 embryos, and the foot and in stomata of developing sporophytes. **D)** *PpCLE4::NGG* signal was detected in sporogenic tissues. **E)** *PpCLE5::NGG* activity was detected in stomata. **F)** *PpCLE6::NGG* was detected in maternal tissue, the sporophyte foot, stomata and a group of cells in the middle of the sporophyte. **G)** *PpCLE7::NGG* signal was detected in maternal tissue, in sporophyte hypophyses, and stomata. **H)** *PpCLE8::NGG* signal was detected in and around stomata, and at the Stage 4 sporophyte tip. **I)** *PpCLE9::NGG* signal was detected in maternal tissue, the sporophyte foot, stomata and a file of cells at the centre of the sporophyte. **J)** *PpCLV1a::NGG* signal was detected in sporogenic tissue**. K)** *PpCLV1b::NGG* activity was present in the apical part of the developing sporophyte. **L)** *PpRPK2::NGG* signal was detected in the basal part and a band in the middle of the sporophyte. Staining times were 3h (*PpCLV1b::NGG*, *PpRPK2::NGG*), 7h (*PpCLE3::NGG*, *PpCLE9::NGG*, *PpCLV1a::NGG*) or 16h (*PpCLE1::NGG*, *PpCLE2::NGG*, *PpCLE4::NGG*, *PpCLE5::NGG*, *PpCLE6::NGG*, *PpCL71::NGG*). Arrowheads indicate stomata with (black) or without (white) staining. Scale bars = 100 µm.

### Different genes are co-expressed in gametophytes and sporophytes

To identify any generalities in our data and further explore potential for co-expression of peptide and receptor encoding gene pairs, we used Pearson’s correlation coefficient to analyse the frequencies of gene expression in all tissues examined in this and previous (Nemec-Venza et al. 2022) work (Figure 7, Supplementary Table 1). Using data from gametophytes, correlation plots showed that amongst peptide-encoding genes, *PpCLE1* and *PpCLE2* had the most similar expression patterns, followed by *PpCLE3* and *PpCLE4*, *PpCLE5* and *PpCLE7* (Figure 7A). Of the receptor-encoding genes, *PpCLV1b* and *PpRPK2* had the most similar expression patterns (Figure 7A). Analysis of sporophyte data showed that *PpCLE4* and *PpCLV1a* expression patterns were most similar, followed by *PpCLE5*, *PpCLE8* and *PpCLE1* (Figure 7B). Of the receptor-encoding genes, *PpCLV1a* and *PpCLV1b* had the most similar expression patterns (Figure 7B). Many genes had highly divergent expression patterns, and this divergence was not detected in gametophyte data. For instance, *PpCLV1a* expression patterns were divergent from *PpCLE1*, *PpCLE3*, *PpCLE5*, *PpCLE7*, *PpCLE8* and *PpCLE9* expression patterns, and the *PpCLV1b* expression pattern was divergent from *PpCLE1*, *PpCLE2*, *PpCLE3* and *PpRPK2* expression patterns (Figure 7B). This analysis showed no strict coupling between peptide and receptor-encoding gene expression patterns and that different genes are co-expressed in gametophytes and sporophytes.

**Figure 7:**
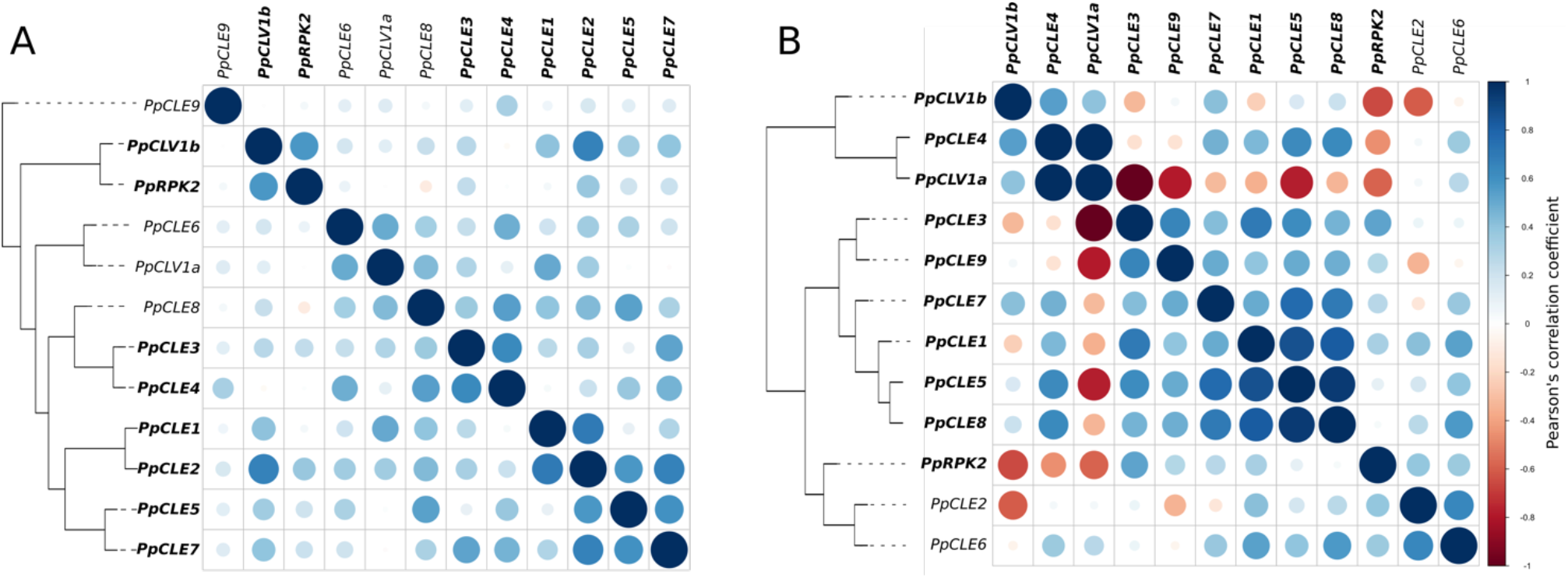
Different *CLAVATA* genes are co-expressed in gametophytes and sporophytes. **A)** Correlation plot showing that *PpCLE1* and *PpCLE2* have the most similar expression patterns in gametophytes, followed by *PpCLE3* and *PpCLE4*, *PpCLE5* and *PpCLE7* and *PpCLV1b* and *PpRPK2*. **B)** Correlation plot showing that *PpCLE4* and *PpCLV1a* have the most similar expression patterns in sporophytes, followed by *PpCLE5*, *PpCLE8* and *PpCLE1*. *PpCLE3* and *PpCLV1a* had the most divergent expression patterns, and *PpCLV1a* and *PpCLV1b* expression patterns were highly divergent from *PpRPK2* expression patterns. Gene names in bold are discussed in the main text. Dendrograms groups are based on Pearson’s correlation coefficient.

## Discussion

### *PpCLE* promoters show cell type specific activities

CLAVATA pathway genes diversified independently from low ancestral copy numbers in mosses and flowering plants (Whitewoods *et al*., 2018). While 32 CLEs and their receptors function in diverse tissues and organs during Arabidopsis development (Cock & McCormick, 2001; Jun *et al*., 2010; Narasimhan & Simon, 2022) and roles for CLAVATA in moss protonema and gametophore development were previously identified (Whitewoods *et al*., 2018; Whitewoods *et al*., 2020; Cammarata *et al*., 2022; Nemec-Venza *et al*., 2022), specific sites of moss CLAVATA activity were previously unknown. Here we used a comprehensive analysis of *PpCLE*, *PpCLV1* and *PpRPK2* promoter activity throughout *P. patens* development to explore such specificity and guide future work. *PpCLE*s are predicted to encode four CLE peptides, RMVPTGPNPLHN (*PpCLE1-3*, *PpCLE8*, *PpCLE9*), RMVPSGPNPLHN (*PpCLE4*), RLVPTGPNPLHN (*PpCLE5*, *PpCLE6*) and RVVPTGPNPLHN (*PpCLE7*). Hence genes encoding the same peptide but expressed in different places could perform similar functions (Table 1, Supplementary Table 1). Promoter activity indicated that PpCLE1-3/8/9 and PpCLE5/6 peptides were likely produced in all tissues, but *PpCLE4* and *PpCLE7* expression marked a subset of tissues with *PpCLE7* showing the narrowest range in gametophytes and *PpCLE4* and *PpCLE7* showing the narrowest range in sporophytes (Table 1, Supplementary Table 1). No *PpCLE* was uniquely expressed in any tissue type, but patterns of promoter activity were unique to individual cell types within tissues, for instance *PpCLE3* promoter activity marked gametophore apical cells and egg cells, *PpCLE5* activity marked rhizoid initiating bud cells, and *PpCLE9* activity marked phyllid midribs. No *PpCLE*s or receptor-encoding genes showed identical expression patterns, and subsets of *PpCLE*s and receptor promoters were active in each cell or tissue type, with no tissues showing no promoter activity (Table 1, Supplementary Table 1). *PpCLV1a*, *PpCLV1b* and *PpRPK2* promoter activities were typically broader than *PpCLE* promoter activities. Overall, our data are consistent with the notion from other species that promoter evolution generates new sites of CLE signalling and might drive divergence of gene function (Hobe *et al*., 2003; Furumizu & Aalen, 2023), providing a framework to identify individual *PpCLE* functions in future work.

**Table 1:** Summary of promoter activities recorded in this study and previous work (Nemec-Venza *et al*., 2022). A) In gametophytes, all promoters were found to be active in caulonema and phyllids, whereas other tissues expressed a subset of *PpCLE*s. While *PpCLE9* was expressed in all tissue classes, other promoter activities were detected in a narrower range of tissues. **B)** In sporophytes, all promoters were found to be active in stomata, whereas other tissues expressed a subset of *PpCLE*s or receptor encoding genes. While *PpCLE6* was expressed in all tissue classes, the promoter activities of other CLAVATA pathway genes were more restricted. Promoter activity was scored as present (+) where it was detected in 50% or more of samples evaluated in a given tissue class; observed frequencies from specific tissue classes are recorded in Supplementary Table 1. No data were recorded in some PpCLV1a tissues due to developmental defects in these lines. *PpCLE*s encoding the same presumptive peptide are shaded using the same colour.

| Tissue | *PpCLE1* | *PpCLE2* | *PpCLE3* | *PpCLE4* | *PpCLE5* | *PpCLE6* | *PpCLE7* | *PpCLE8* | *PpCLE9* | *PpCLV1a* | *PpCLV1b* | *PpRPK2* |
| --- | --- | --- | --- | --- | --- | --- | --- | --- | --- | --- | --- | --- |
| Chloronema |  |  |  | + |  |  |  | + | + | + |  | + |
| Caulonema | + | + | + | + | + | + | + | + | + | + | + | + |
| Gametophore buds |  | + | + | + | + | + | + |  | + |  | + | + |
| Gametophore apices |  |  | + | + |  | + | + |  | + |  | + | + |
| Gametophore axes |  | + |  |  | + | + |  |  | + | + | + | + |
| Phyllids | + | + | + | + | + | + | + | + | + | + | + | + |
| Gametophore rhizoids |  |  |  | + | + |  |  | + | + | + | + |  |
| Antheridia |  |  |  |  |  | + |  |  | + | + |  | + |
| Archegonia and eggs |  |  | + | + |  | + |  |  | + | + | + | + |

B) Sporophytic tissues
| Tissue | Stage | *PpCLE1* | *PpCLE2* | *PpCLE3* | *PpCLE4* | *PpCLE5* | *PpCLE6* | *PpCLE7* | *PpCLE8* | *PpCLE9* | *PpCLV1a* | *PpCLV1b* | *PpRPK2* |
| --- | --- | --- | --- | --- | --- | --- | --- | --- | --- | --- | --- | --- | --- |
| Apical embryo | 1 to 2 |  |  | + |  |  | + |  |  | + |  | + |  |
| Basal embryo | 1 to 2 |  | + | + |  |  | + |  |  |  |  |  | + |
| Sporophyte foot | 3 to 7 | + | + | + |  |  | + |  |  | + | No data |  | + |
| Intercalary region | 4 to 5 |  | + | + |  |  | + | + |  | + |  | + | + |
| Sporogenic tissue | 4 to 5 |  | + |  | + |  | + |  |  |  | + | + |  |
| Stomata | 6 to 7 | + | + | + | + | + | + | + | + | + | No data | + | + |
| Capsule | 7 |  | + | + |  |  | + |  | + | + | No data | + |  |

### Gametophytes and sporophytes show specialised CLAVATA activities

Divergence in CLAVATA expression patterns between gametophytes and sporophytes appeared a general feature in our previously published (Nemec-Venza *et al*., 2022) and new data (Figure 7, Table 1). In gametophytes, all components of the *P. patens* CLAVATA pathway were active in caulonemata and phyllids, and these expression patterns are consistent with caulonemal and leaf defects in mutants (Whitewoods *et al*., 2018; Whitewoods *et al*., 2020; Nemec-Venza *et al*., 2022). In sporophytes, all promoters were active in stomata, by comparison with Arabidopsis stomatal development (Qian *et al*., 2018) indicating a potential ancestral role for CLE signalling. Whilst most gametophytic tissues showed significant overlap of *PpCLV1b* and *PpRPK2* promoter activities, promoter activities marked clearly distinct apical (*PpCLV1b::NGG*) and basal (*PpRPK2::NGG*) embryonic domains, thus diverging between generations (Figure 1, Figure 6, Figure 7, Table 1). *PpCLE* promoter activities also differed between generations (Table 1). For instance, the *PpCLE1* promoter was active in a narrow range of tissues in gametophytes and sporophytes, but *PpCLE4*, *PpCLE5, PpCLE7* and *PpCLE8* were expressed more broadly in gametophytes than sporophytes (Table 1). Our data suggest that CLAVATA functions may have diversified in response to distinct selection pressures acting on gametophytes and sporophytes, perhaps accounting for high gene copy numbers in *P. patens* relative to other bryophytes (Whitewoods *et al*., 2018).

### Sites of receptor expression overlap with auxin sensing minima in multiple developmental contexts

Whilst *PpRPK2* acts in parallel with cytokinin to regulate gametophore bud development, and independently from *PpCLV1* (Cammarata *et al*., 2022), *PpRPK2* and *PpCLV1* both regulate auxin synthesis and auxin transporter (*PpPINs A-C*) expression to determine plant spread in protonemata (Nemec-Venza *et al*., 2022). The data presented here identify caulonemal apical cells, gametophore apical cells, the base of developing leaves, rhizoid tips and pre-egg and egg cells as regions of high *PpRPK2* and *PpCLV1* promoter activity, coinciding with previously identified sites of minimal auxin sensing (Thelander *et al*., 2019; Landberg *et al*., 2021). Moreover, *PpPIN* expression domains are similar to CLAVATA expression domains in leaves and reproductive tissues (Landberg *et al*., 2013; Viaene *et al*., 2014; Lüth *et al*., 2023), and *Pppin* mutants have fertility defects (Bennett *et al*., 2014; Lüth *et al*., 2023), a phenotype shared by CLAVATA receptor mutants (Figures 3-5). Auxin synthesis is required for normal egg development, and biosynthetic *Ppshi*-*2* mutant eggs resemble *Pprpk2* mutant eggs (Landberg *et al*., 2013). These data suggest that CLAVATA receptor signalling could modulate auxin homeostasis in multiple developmental contexts in *P. patens*, and divergent receptor mutant phenotypes show that *PpRPK2* and *PpCLV1* act independently in reproductive development as well as vegetative development (Cammarata *et al*., 2022).

### CLAVATA activity is divergent in gametophyte and sporophyte meristems

The Arabidopsis CLAVATA pathway is best known for its activity in regulating the size of the meristematic stem cell pool (Clark *et al*., 1997), yet few homologies in gross meristem structure or meristematic gene functions have been identified within land plants (Fouracre & Harrison, 2022). Whilst moss gametophyte meristems comprise a single apical cell, embryonic apical cell and later intercalary proliferation are separated in time and space by sporangium development in sporophytes (Parihar, 1959). Juxtaposition of persistently active apical stem cells with an intercalary proliferative region is proposed to have enabled the elaboration of sporophyte growth during vascular plant evolution (Harrison, 2017). The moss KNOX protein MKN2 promotes such intercalary proliferation by upregulating cytokinin biosynthesis (Singer & Ashton, 2007; Sakakibara *et al*., 2008; Coudert *et al*., 2019), and shared KNOX functions in vascular plants indicate potential homology in mechanisms for meristematic proliferation (Harrison *et al*., 2005; Jasinski *et al*., 2005; Yanai *et al*., 2005). *PpCLV1b* and *PpRPK2* promoter activities abut or overlap at the site of the intercalary meristem where multiple CLEs are also produced, bringing potential for interaction with MKN2 and/or cytokinin in the regulation of meristematic cell proliferation. Our data are therefore consistent with a model in which the moss intercalary meristem shares homology with the vascular plant shoot apical meristem as they are both regulated by genetic networks that were present in the ancestral land plant. However, *clavata* mutant infertility currently makes this model difficult to test. Given that we detected no *PpCLE* expression specific to embryonic apical cells, our data also support the notion that the genetic networks for stem cell function are divergent in gametophytes and sporophytes (Frank & Scanlon, 2015; Harrison, 2015).

## Materials and methods

### Plant material and growth conditions

The Gransden strain of *P. patens* was used as the wild-type (WT) strain for all experiments, and *Ppclv1a*, *Ppclv1b*, *Ppclv1a1b*, *Pprpk2*, *Ppclv1a1brpk2*, *PpCLE1::NGG*, *PpCLE2::NGG*, *PpCLE3::NGG, PpCLE4::NGG, PpCLE5::NGG, PpCLE6::NGG, PpCLE7::NGG*, *PpCLE8::NGG, PpCLE9::NGG, PpCLV1A::NGG*, *PpCLV1B::NGG* and *PpRPK2::NGG* and mutant line generation strategies were previously described (Whitewoods *et al*., 2018; Whitewoods *et al*., 2020; Cammarata *et al*., 2022; Nemec-Venza *et al*., 2022). With the exception of *PpCLV1A::NGG,* where expression was analysed in 3 lines due to sporophyte development defects, a single reporter line was used for each gene.

For vegetative expression analyses, plants were spot propagated on BCDAT media containing 0.5% agar as in a previous study (Nemec-Venza *et al*., 2022) and grown at 23°C in continuous light or at 22°C in long day conditions (16h light/8h dark). To induce sexual reproduction, protonemal homogenates were propagated in Magenta vessels containing hydrated peat plugs with 30 ml half-strength BCD media and grown at 23 °C in continuous light for 8–10 weeks prior to transfer to 16 °C short day conditions (16h dark/8h light). To stain gametangia and phenotype plants, tissue was harvested 2–3 weeks after induction. At this point, sterile water was repeatedly poured on the tissue until it was wet enough to allow fertilization. To stain young and mature sporophytes from *promoter::NGG* lines, tissue was respectively collected 12 and 24 days after fertilization. For the receptor mutant lines, cultures were maintained for 2 months following fertilization to determine sporophyte production rates.

### Histochemical analysis and imaging

GUS staining was performed at 37°C with a solution containing 100 mM phosphate buffer (pH 7.0), 10 mM Tris-HCl (pH 8.0), 1 mM ethylenediaminetetraacetic acid (EDTA), 0.05% Triton X-100, 2 mM potassium ferricyanide, 2 mM potassium ferrocyanide and 1 mg/ml X-Gluc (5-bromo-4-chloro-3-indolyl-β-d-glucuronic acid) dissolved in 10% (v/v) DMSO. Staining times for buds and leaves were 7.5 h for *PpCLE3::NGG*, *PpCLE4::NGG*, *PpCLE5::NGG*, *PpCLE6::NGG*, *PpCLV1b::NGG* and *PpRPK2::NGG*, 15 h for *PpCLE2::NGG*, *PpCLE8::NGG*, *PpCLE9::NGG* and *PpCLV1a::NGG* and 21 h for *PpCLE1::NGG* and *PpCLE7::NGG*. Antheridia, archegonia and sporophyte staining times are indicated in figures or figure legends. After staining, samples were cleared in 70% ethanol for 24h. Imaging was performed using a Keyence VHX-1000 digital microscope (whole gametophores and leaves), a Leica DM2000 microscope (buds) or a Zeiss AxioImager.M2 microscope (antheridia, archegonia and sporophytes).

### Mutant phenotyping

Wild-type, *Ppclv1a*, *Ppclv1b*, *Pprpk2* and *Ppclv1aclv1brpk2* mutants were grown in reproduction inducing conditions. Gametangia (antheridia and archegonia) were dissected from the fresh apices and quantification of gametangium morphology was performed using at least 2 experimental replicates. To assess fertility, sporophytes from 3 experimental replicates were counted two months after fertilization. A GXML 1500 compound microscope was used to image tissues and the data were subsequently analysed in ImageJ. Statistical analysis was performed using R Statistical Software (v4.1.2) (R Core Team, 2021).

### Sperm cell imaging

To visualize sperm cells, whole gametophores were cleared and mounted in Hoyer’s solution (50 ml distilled water, 30 g Gum Arabic, 200 g Chloral Hydrate, 20 ml Glycerin) and rinsed in water. Antheridia were then dissected and stained with 1mg/L DAPI as detailed in (Horst & Reski, 2017). Imaging was performed with a Zeiss AxioImager.M2 epifluorescence microscope.

### Correlation analysis

Dendrograms were based on Pearson’s correlation coefficient as calculated by cor() and generated based on Euclidean distance with the hclust() function using the “average” method with R Statistical Software (v4.1.2) (R Core Team, 2021). The corrplot() function was used to generate correlation plots.

## Acknowledgements

We thank the Gatsby Charitable Foundation, the Bristol Centre for Agricultural Innovation and BBSRC for funding. We thank Wei Liu and Yasuko Kamisugi for technical help. We thank Mauro Tretiach for hosting a visit from ZNV. We thank Jim Fouracre and Adrienne Roeder for commenting on a manuscript draft. CJH thanks Lara Gibbs and her NHS colleagues for making this work possible.

## Conflicts of interest

The authors declare no conflicting interests.

## Author contributions

All authors contributed to design of the study, and ZNV and GG undertook the experimental work and data analyses with supervision from CJH. All authors contributed to manuscript preparation and editing.

## Supplementary data

**Supplementary Figure 1:**
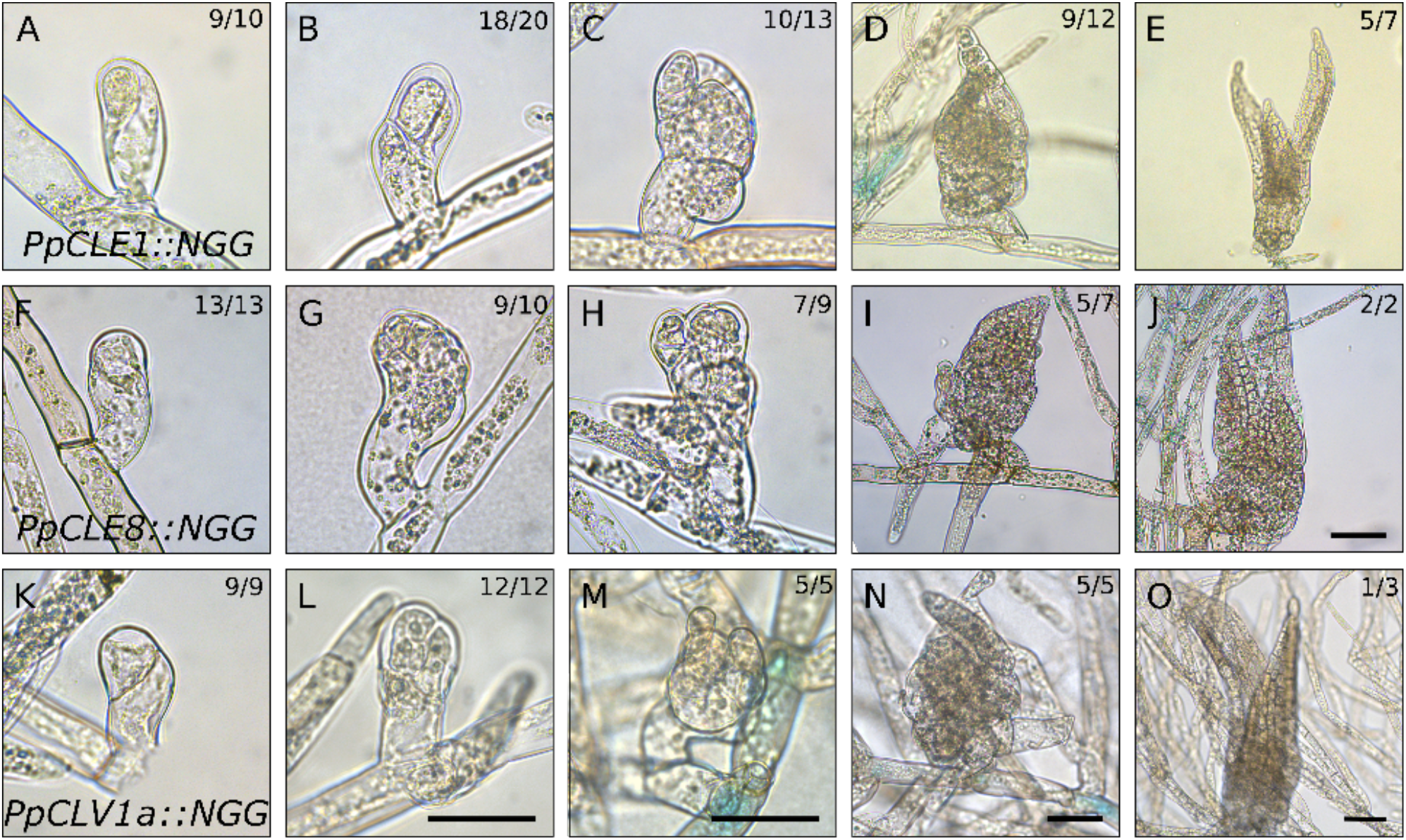
Absent *CLAVATA* promote activity in gametophore initiation. **(A-E)** *PpCLE1::NGG* was usually inactive at **(A)** Stage 0, **(B)** Stage 1, **(C)** Stage 2, **(D)** Stage 3 and **(E)** Stage 4 of gametophore development. **(F-J)** *PpCLE8::NGG* was usually inactive at **(F)** Stage 0, **(G)** Stage 1, **(H)** Stage 2, **(I)** Stage 3 and **(J)** Stage 4 of gametophore development. **(K-O)** *PpCLV1a::NGG* was usually inactive at **(K)** Stage 0, **(L)** Stage 1, **(M)** Stage 2, **(N)** Stage 3 and **(O)** Stage 4 of gametophore development. Numbers indicate the proportion of buds with a similar expression pattern. Scale bars Stages 0-2 = 50 µm, Scale bars at Stages 3-4 = 100 µm.

**Supplementary Figure 2:**
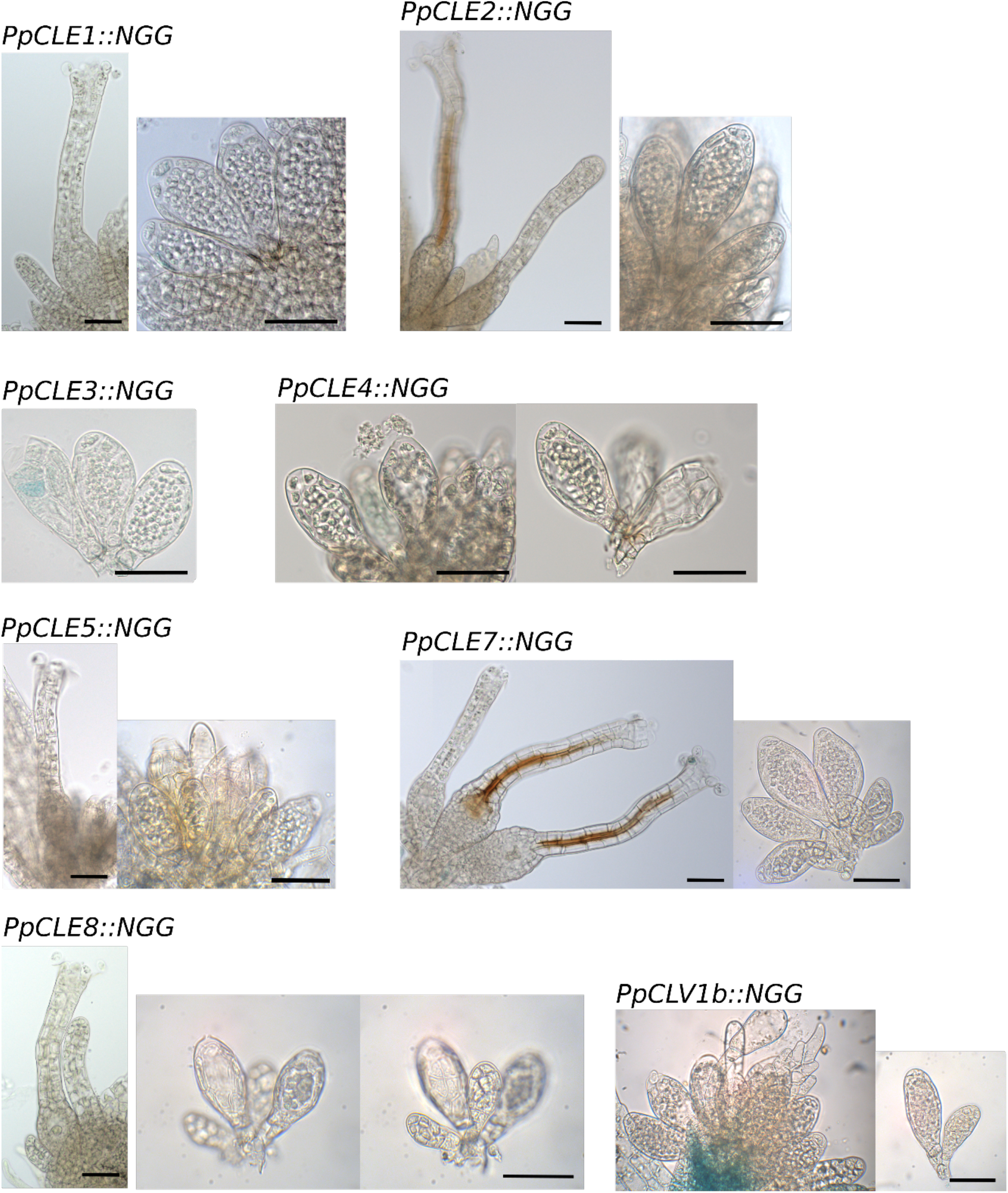
GUS-stained gametangia from lines showing low or no expression. Less than 50% of antheridia in *PpCLE1::NGG*, *PpCLE2::NGG*, *PpCLE3::NGG*, *PpCLE4::NGG*, *PpCLE5::NGG*, *PpCLE7::NGG*, *PpCLE8::NGG* and *PpCLV1b::NGG* samples showed expression. Less than 50% of archegonia from *PpCLE1::NGG*, *PpCLE2::NGG*, *PpCLE5::NGG*, *PpCLE7::NGG* and *PpCLE8::NGG* lines showed signal. Scale bar = 50 µm

**Supplementary Table 1:** Promoter activities recorded in this study and previous work. (Nemec-Venza *et al*., 2022). The frequency that signal was detected in specified cells and tissue types is shown. Where the frequency was 50% or more, promoter activity was scored as present for Table 1 shown in the main text, and bold lines delimit tissue classes used in the summary table. No data were recorded in some *PpCLV1a::NGG* tissues due to developmental defects in these lines. *PpCLE*s encoding the same presumptive peptide are shaded using the same colour.

| Tissue class for Table 1 | Classes scored for similarity plot | PpCLE1 | PpCLE2 | PpCLE3 | PpCLE4 | PpCLE5 | PpCLE6 | PpCLE7 | PpCLE8 | PpCLE9 | PpCLV1a | PpCLV1b | PpRPK2 |
| --- | --- | --- | --- | --- | --- | --- | --- | --- | --- | --- | --- | --- | --- |
| Chloronema | Germinating spore | 0.00 | 0.03 | 0.14 | 0.00 | 0.02 | 0.09 | 0.03 | 0.10 | 0.09 | **0.58** | 0.03 | 0.03 |
|  | Primary chloronema | 0.30 | 0.00 | 0.13 | 0.27 | 0.10 | 0.21 | 0.08 | **0.50** | **0.63** | **1.00** | 0.11 | 0.19 |
|  | Chloronema branches | 0.13 | 0.27 | 0.33 | **0.50** | 0.46 | 0.31 | 0.46 | 0.00 | **0.70** | 0.11 | 0.00 | **0.79** |
| Caulonema | Primary caulonema | **0.53** | **0.69** | **0.80** | **0.83** | **0.87** | **0.86** | **1.00** | **0.63** | **0.88** | **1.00** | **1.00** | **0.97** |
|  | Caulonema tips | 0.09 | **0.76** | 0.46 | **0.52** | **0.94** | **0.51** | **0.91** | 0.19 | **0.91** | 0.12 | **0.63** | **1.00** |
| Gametophore buds | Cell under the bud (Stage 0-1) | 0 | 0.21 | 0.08 | 0.27 | **0.58** | **0.8** | **0.68** | 0.09 | 0.32 | 0.09 | **0.57** | **0.73** |
|  | Stage 0 | 0.10 | 0.00 | 0.06 | 0.00 | 0.10 | 0.00 | 0.20 | 0.00 | **0.67** | 0.00 | **0.63** | **0.92** |
|  | Stage 1 apical | 0.13 | 0.00 | **0.59** | 0.24 | 0.06 | 0.00 | 0.21 | 0.10 | 0.31 | 0.00 | 0.43 | **0.74** |
|  | Stage 1 basal | 0.00 | 0.27 | 0.00 | 0.05 | 0.25 | 0.00 | 0.21 | 0.00 | 0.13 | 0.00 | **0.70** | **0.63** |
|  | Stage 2-4 apex | 0.00 | 0.00 | **1.00** | **0.89** | 0.00 | 0.35 | **0.88** | 0.32 | **0.53** | 0.00 | 0.00 | 0.00 |
|  | Stage 2-4 broader apex | 0.25 | 0.09 | 0.00 | 0.00 | 0.13 | 0.00 | 0.00 | 0.00 | 0.00 | 0.00 | **0.93** | **0.86** |
|  | Stage 2-4 rhizoid | 0.44 | **0.59** | 0.07 | 0.28 | **0.87** | 0.38 | **0.83** | 0.21 | 0.47 | 0.00 | 0.29 | **0.71** |
| Gametophore apex | Gametophore apex | 0.00 | 0.44 | **1.00** | **1.00** | 0.37 | **1.00** | **0.90** | 0.05 | **1.00** | 0.38 | **0.85** | **1.00** |
| Gametophore axis | Gametophore axis | 0.04 | **0.59** | 0.06 | 0.03 | **0.50** | **1.00** | 0.24 | 0.27 | **0.93** | **1.00** | **1.00** | **0.85** |
| Phyllids | Phyllid lamina | **1.00** | **1.00** | **1.00** | 0.00 | 0.00 | 0.00 | **1.00** | 0.00 | **0.50** | **1.00** | **1.00** | **1.00** |
|  | Phyllid base | **1.00** | **1.00** | **1.00** | **1.00** | **0.50** | **1.00** | **0.50** | **1.00** | **0.50** | **1.00** | **1.00** | **1.00** |
|  | Phyllid midrib | **1.00** | **0.50** | 0.00 | 0.00 | 0.00 | **1.00** | 0.00 | 0.00 | **1.00** | **1.00** | **0.50** | 0.20 |
| Gametophore rhizoids | Gametophore rhizoids | 0.11 | 0.44 | 0.32 | **0.56** | **1.00** | 0.41 | 0.38 | **0.64** | **0.52** | **0.52** | **0.52** | 0.41 |
| Antheridia | Antheridia (Stage 1-5 apex) | 0.00 | 0.00 | 0.00 | 0.19 | 0.00 | 0.00 | 0.00 | 0.03 | **1.00** | **0.53** | 0.13 | **0.98** |
|  | Antheridia (Stage 1-5 base) | 0.00 | 0.00 | 0.42 | 0.19 | 0.00 | **1.00** | 0.00 | 0.03 | 0.00 | **0.97** | 0.13 | **0.98** |
|  | Antheridia (Stage 6-8 apex) | 0.24 | 0.02 | 0.44 | 0.41 | 0.29 | 0.00 | 0.20 | 0.30 | **1.00** | 0.40 | 0.04 | 0.30 |
|  | Antheridia (Stage 6-8 base) | 0.24 | 0.02 | 0.44 | 0.41 | 0.29 | **1.00** | 0.20 | 0.30 | 0.00 | **0.90** | 0.04 | 0.30 |
| Archegonia and eggs | Egg and canal cells (Stage 4-8) | 0.00 | 0.00 | **0.96** | **0.77** | 0.14 | **0.64** | 0.00 | 0.10 | **0.95** | **1.00** | 0.07 | **0.91** |
|  | Eggs at (Stage 9-10) | 0.00 | 0.00 | **0.75** | 0.10 | 0.08 | 0.20 | 0.00 | 0.00 | **0.62** | 0.00 | **0.90** | **1.00** |
|  | Archegonium tip | 0.00 | 0.00 | 0.00 | 0.00 | 0.00 | 0.00 | 0.00 | 0.00 | **1.00** | 0.25 | 0.00 | **0.50** |
|  | Archegonium neck (Stage 7-9) | 0.00 | 0.00 | 0.00 | **0.77** | 0.00 | **0.71** | 0.00 | 0.00 | **1.00** | 0.00 | 0.00 | 0.30 |
|  | Archegonium venter | 0.00 | 0.15 | 0.00 | 0.10 | 0.00 | 0.30 | 0.00 | 0.00 | 0.10 | 0.00 | 0.20 | **0.97** |
| Sporophyte development | Maternal tissue | 0.46 | **1.00** | **0.63** | 0.00 | 0.27 | **1.00** | 0.14 | 0.40 | **1.00** | 0.00 | 0.00 | **1.00** |
| Apical embryo | Apical embryo (Stage 1-2) | 0.00 | 0.00 | **0.67** | 0.00 | 0.09 | **0.50** | 0.00 | 0.00 | **1.00** | 0.00 | **1.00** | 0.00 |
| Basal embryo | Basal embryo (Stage 1-2) | 0.18 | **1.00** | **0.67** | 0.00 | 0.18 | **0.67** | 0.00 | 0.00 | 0.33 | 0.00 | 0.00 | **0.67** |
| Sporophyte foot | Sporophyte foot (Stage 3-7) | **0.60** | **0.96** | **0.95** | 0.00 | 0.21 | **1.00** | 0.00 | 0.14 | **0.52** | No data | 0.00 | **1.00** |
| Intercalary region | Intercalary region (Stage 4-5) | 0.00 | **0.50** | **0.63** | 0.00 | 0.17 | **0.93** | **0.67** | 0.13 | **0.77** | 0.00 | **1.00** | **1.00** |
| Sporogenic tissue | Sporogenic tissue (Stage 3-5) | 0.00 | **0.67** | 0.00 | **0.57** | 0.00 | **0.90** | 0.00 | 0.00 | 0.00 | **1.00** | **1.00** | 0.00 |
| Stomata | Stomata (Stage 6-7) | **1.00** | **0.71** | **1.00** | **0.80** | **1.00** | **1.00** | **0.92** | **1.00** | **1.00** | No data | **1.00** | **0.50** |
| Capsule | Capsule (Stage 7) | 0.25 | **1.00** | **0.50** | 0.30 | 0.31 | **1.00** | 0.08 | **0.50** | **0.50** | No data | **1.00** | 0.00 |

## Notes

### Competing Interest Statement

The authors have declared no competing interest.

## References

Aoyama T, Hiwatashi Y, Shigyo M, Kofuji R, Kubo M, Ito M, Hasebe M. 2012. AP2-type transcription factors determine stem cell identity in the moss *Physcomitrella patens*. Development 139: 3120–3129.

Bennett TA, Liu MM, Aoyama T, Bierfreund NM, Braun M, Coudert Y, Dennis RJ, O’Connor D, Wang XY, White CD, Decker EL, Reski R, Harrison CJ. 2014. Plasma membrane-targeted PIN proteins drive shoot development in a moss. Current Biology 24: 2776–2785.

Brand U, Fletcher JC, Hobe M, Meyerowitz EM, Simon R. 2000. Dependence of stem cell fate in Arabidopsis on a feedback loop regulated by CLV3 activity. Science 289: 617–619.

Cammarata J, Farfan CM, Scanlon MJ, Roeder AHK. 2022. Cytokinin–CLAVATA cross-talk is an ancient mechanism regulating shoot meristem homeostasis in land plants. PNAS 119: e2116860119.

Clark SE, Running MP, Meyerowitz EM. 1993. *Clavata1*, a regulator of meristem and flower development in *Arabidospis*. Development 119: 397–418.

Clark SE, Williams RW, Meyerowitz EM. 1997. The Clavata1 gene encodes a putative receptor kinase that controls shoot and floral meristem size in Arabidopsis. Cell 89: 575–585.

Cock JM, McCormick S. 2001. A large family of genes that share homology with CLAVATA3. Plant Physiology 126: 939–942.

Coudert Y, Novák O, Harrison CJ. 2019. A KNOX-cytokinin regulatory module predates the origin of indeterminate vascular plants. Current Biology 29: 2743–2750.

Fletcher JC. 2018. The CLV-WUS stem cell signaling pathway: a roadmap to crop yield optimization. Plants 7: 87.

Fletcher JC. 2020. Recent Advances in Arabidopsis CLE Peptide Signaling. Trends in Plant Science 25: 1005–1016.

Fletcher JC, Brand U, Running MP, Simon R, Meyerowitz EM. 1999. Signalling of cell fate decisions by CLAVATA3 in Arabidopsis shoot meristems. Science 283: 1911–1914.

Fouracre JP, Harrison CJ. 2022. How was apical growth regulated in the ancestral land plant? Insights from the development of non-seed plants. Plant Physiol 190: 100–112.

Frank MH, Scanlon MJ. 2015. Cell-specific transcriptomic analyses of three-dimensional shoot development in the moss *Physcomitrella patens*. Plant Journal 83: 743–751.

Furumizu C, Aalen RB. 2023. Peptide signaling through leucine-rich repeat receptor kinases: insight into land plant evolution. New Phytologist 238: 977–982.

Goad DM, Zhu C, Kellogg EA. 2017. Comprehensive identification and clustering of CLV3/ESR-related (CLE) genes in plants finds groups with potentially shared function. New Phytologist 216: 605–616.

Harrison CJ. 2015. Shooting through time: new insights from transcriptomic data. Trends in Plant Science 20: 468–470.

Harrison CJ. 2017. Development and genetics in the evolution of land plant body plans. Philosophical Transactions of the Royal Society B 372: e20150490.

Harrison CJ, Corley SB, Moylan EC, Alexander DL, Scotland RW, Langdale JA. 2005. Independent recruitment of a conserved developmental mechanism during leaf evolution. Nature 434: 509–514.

Hirakawa Y, Uchida N, Yamaguchi YL, Tabata R, Ishida S, Ishizaki K, Nishihama R, Kohchi T, Sawa S, Bowman JL. 2019. Control of proliferation in the haploid meristem by CLE peptide signalling in *Marchantia polymorpha*. PLOS Genetics 15: e1007997.

Hirakawa Y, Fujimoto T, Ishida S, Uchida N, Sawa S, Kiyosue T, Ishizaki K, Nishihama R, Kohchi T, Bowman JL. 2020. Induction of multichotomous branching by CLAVATA peptide in *Marchantia polymorpha*. Current Biology 30: 3833–3840 e3834.

Hobe M, Muller R, Grunewald M, Brand U, Simon R. 2003. Loss of CLE40, a protein functionally equivalent to the stem cell restricting signal CLV3, enhances root waving in Arabidopsis. Development Genes and Evolution 213: 371–381.

Horst NA, Reski R. 2017. Microscopy of Physcomitrella patens sperm cells.. Plant Methods 13.

Jasinski S, Piazza P, Craft J, Hay A, Woolley L, Rieu I, Phillips A, Hedden P, Tsiantis M. 2005. KNOX action in Arabidopsis is mediated by coordinate regulation of cytokinin and gibberellin activities. Curr Biol 15: 1560–1565.

Jun J, Fiume E, Roeder AH, Meng L, Sharma VK, Osmont KS, Baker C, Ha CM, Meyerowitz EM, Feldman LJ, et al. 2010. Comprehensive analysis of CLE polypeptide signaling gene expression and overexpression activity in Arabidopsis. Plant Physiology 154: 1721–1736.

Kinoshita A, Betsuyaku S, Osakabe Y, Mizuno S, Nagawa S, Stahl Y, Simon R, Yamaguchi-Shinozaki K, Fukuda H, Sawa S. 2010. RPK2 is an essential receptor-like kinase that transmits the CLV3 signal in Arabidopsis. Development 137: 3911–3920.

Kofuji R, Hasebe M. 2014. Eight types of stem cells in the life cycle of the moss *Physcomitrella patens*. Current Opinion in Plant Biology 17: 13–21.

Landberg K, Pederson ER, Viaene T, Bozorg B, Friml J, Jönsson H, Thelander M, Sundberg E. 2013. The moss Physcomitrella patens reproductive organ development is highly organized, affected by the two SHI/STY genes and by the level of active auxin in the SHI/STY expression domain. Plant Physiology 162: 1406–1419.

Landberg K, Šimura J, Ljung K, Sundberg E, Thelander M. 2021. Studies of moss reproductive development indicate that auxin biosynthesis in apical stem cells may constitute an ancestral function for focal growth control. New Phytologist 229: 845–860.

Leyser HMO, Furner IJ. 1992. Characterization of three shoot apical meristem mutants of *Arabidopsis thaliana*. Development 116: 397–403.

Lüth VM, Rempfer C, van Gessel N, Herzog O, Hanser M, Braun M, Decker EL, Reski R. 2023. A Physcomitrella PIN acts in spermatogenesis and sporophyte retention. New Phytologist 237: 2118–2135.

Mayer KFX, Schoof H, Haecker A, Lenhard M, Jurgens G, Laux T. 1998. Role of Wuschel in regulating stem cell fate in the Arabidopsis shoot meristem. Cell 95: 805–815.

Morris JL, Puttick MN, Clark JW, Edwards D, Kenrick P, Pressel S, Wellman CH, Yang Z, Schneider H, Donoghue PCJ. 2018. The timescale of early land plant evolution. PNAS 115: E2274–E2283.

Narasimhan M, Simon R. 2022. Spatial range, temporal span, and promiscuity of CLE-RLK signaling. Frontiers in Plant Science 13: 906087.

Nemec-Venza Z, Madden C, Stewart A, Liu W, Novák O, Pencik A, Cuming AC, Kamisugi Y, Harrison CJ. 2022. CLAVATA modulates auxin homeostasis and transport to regulate stem cell identity and plant shape in a moss. New Phytologist 234: 149–163.

Parihar NS. 1959. Bryophyta: Central Book Depot.

Qian P, Song W, Yokoo T, Minobe A, Wang G, Ishida T, Sawa S, Chai J, Kakimoto T. 2018. The CLE9/10 secretory peptide regulates stomatal and vascular development through distinct receptors. Nature Plants 4: 1071–1081.

Rojo E, Sharma VK, Kovalev V, Raikhel NV, Fletcher JC. 2002. CLV3 is localized to the extracellular space, where it activates the Arabidopsis CLAVATA stem cell signaling pathway. Plant Cell 14: 969–977.

Sakakibara K, Nishiyama T, Deguchi H, Hasebe M. 2008. Class 1 *KNOX* genes are not involved in shoot development in the moss *Physcomitrella patens* but do function in sporophyte development. Evolution and Development 10: 555–566.

Sakakibara K, Reisewitz P, Aoyama T, Friedrich T, Ando S, Sato Y, Tamada Y, Nishiyama T, Hiwatashi Y, Kurata T, et al. 2014. *WOX13*-like genes are required for reprogramming of leaf and protoplast cells into stem cells in the moss *Physcomitrella patens*. Development 141: 1660–1670.

Schoof H, Lenhard M, Haecker A, Mayer KF, Jurgens G, Laux T. 2000. The stem cell population of Arabidopsis shoot meristems in maintained by a regulatory loop between the CLAVATA and WUSCHEL genes [In Process Citation]. Cell 100: 635–644.

Singer SD, Ashton NW. 2007. Revelation of ancestral roles of KNOX genes by a functional analysis of Physcomitrella homologues. Plant Cell Reports 26: 2039–2054.

R Core Team. 2021. R: A language and environment for statistical computing.

Thelander M, Landberg K, Sundberg E. 2019. Minimal auxin sensing levels in vegetative moss stem cells revealed by a ratiometric reporter. New Phytologist 224: 775–788.

Trotochaud AE, Jeong S, Clark SE. 2000. CLAVATA3, a multimeric ligand for the CLAVATA1 receptor-kinase. Science 289: 613–617.

Tvorogova VE, Krasnoperova EY, Potsenkovskaia EA, Kudriashov AA, Dodueva IE, Lutova LA. 2021. What does the WOX say? Review of regulators, targets, partners. Molecular Biology 55: 311–337.

Viaene T, Landberg K, Thelander M, Medvecka E, Pederson E, Feraru E, Cooper ED, Karimi M, Delwiche CF, Ljung K, et al. 2014. Directional auxin transport mechanisms in early diverging land plants. Current Biology 24: 2786–2791.

Whitewoods CD, Cammarata J, Nemec Venza Z, Sang S, Crook AD, Aoyama T, Wang XY, Waller M, Kamisugi Y, Cuming AC, et al. 2018. CLAVATA was a genetic novelty for the morphological innovation of 3D growth in land plants. Current Biology 28: 2365–2376 e2365.

Whitewoods CD, Cammarata J, Nemec Venza Z, Sang S, Crook AD, Aoyama T, Wang XY, Waller M, Kamisugi Y, Cuming AC, et al. 2020. CLAVATA was a genetic novelty for the morphological innovation of 3D growth in land plants. Current Biology 30: 2645–2648.

Willoughby AC, Nimchuk ZL. 2021. WOX going on: CLE peptides in plant development. Current Opinion in Plant Biology 63: 102056.

Wu CC, Li FW, Kramer EM. 2019. Large-scale phylogenomic analysis suggests three ancient superclades of the WUSCHEL-RELATED HOMEOBOX transcription factor family in plants. PLoS One 14: e0223521.

Yanai O, Shani E, Dolezal K, Tarkowski P, Sablowski R, Sandberg G, Samach A, Ori N. 2005. *Arabidopsis* KNOXI proteins activate cytokinin biosynthesis. Current Biology 15: 1566–1571.

